# Activation of endoplasmic reticulum stress via clustering of the inner nuclear membrane protein SUN2

**DOI:** 10.1101/2021.09.14.460295

**Authors:** Sandra Vidak, Leonid A. Serebryannyy, Gianluca Pegoraro, Tom Misteli

## Abstract

One of the major cellular mechanisms to ensure protein homeostasis is the endoplasmic reticulum (ER) stress response. This pathway is typically triggered by accumulation of misfolded proteins in the ER lumen. Here we describe activation of ER stress via protein aggregation in the cell nucleus. We find in the premature aging disease Hutchinson-Gilford Progeria Syndrome (HGPS) activation of ER stress due to the aggregation of the diseases-causing progerin protein at the nuclear envelope. The presence of nucleoplasmic protein aggregates is sensed and signaled to the ER lumen via immobilization and clustering of the inner nuclear membrane protein SUN2, leading to activation of the Unfolded Protein Response (UPR). These results identify a nuclear trigger of ER stress and they provide insight into the molecular disease mechanisms of HGPS.

## INTRODUCTION

The cellular levels of proteins and their homeostasis in response to environmental cues are determined by a finely tuned balance between protein synthesis and degradation. In addition to regulated synthesis of proteins, several major cellular pathways are involved in the maintenance of the proteome, including the proteasomal system and autophagy, which are responsible for removal of damaged, misfolded and aggregated proteins from cells (Martinez et al., 2017). Furthermore, to ensure proper folding of nascent proteins and to prevent the accumulation of disease-causing aggregates, cells employ a repertoire of chaperones that resolubilize protein aggregates and assist in refolding of misfolded or unfolded proteins (Ciechanover and Kwon, 2017; Saibil, 2013).

A prominent cellular mechanism of protein quality control involves the endoplasmic reticulum (ER), the site of protein synthesis and folding of secreted and membrane-bound proteins (Araki and Nagata, 2012). Stringent quality control systems in the lumen of the ER ensure that only correctly folded and functional proteins are released into the secretory pathway and that misfolded proteins are degraded (Zhang and Ye, 2014). Proper functioning of the ER and its protein quality control mechanisms are essential for cellular activities and cell survival. Stimuli that interfere with ER function, such as hypoxia, aberrant calcium homeostasis, viral infections, or overproduction of secreted proteins, disrupt ER homeostasis and cause accumulation of unfolded or misfolded proteins in the ER lumen, resulting in ER stress (Hetz, 2012; Lafleur et al., 2013; Walter and Ron, 2011). ER transmembrane proteins, including inositol-requiring protein 1 (IRE1), PKR-like endoplasmic reticulum kinase (PERK), and activating transcription factor (ATF)-6 detect the onset of ER stress and initiate the Unfolded Protein Response (UPR) signaling pathway to restore normal ER functions (Kozutsumi et al., 1988; Lin et al., 2008; Wang and Kaufman, 2016). Under physiological conditions, the activation of these sensors is inhibited by the binding of their luminal domains by the main ER-resident chaperone GRP78/BiP (78-kDa glucose-regulated protein). Due to its higher affinity to unfolded proteins compared to ER stress sensors, accumulated unfolded proteins cause the dissociation of GRP78/BiP from ER stress sensors and subsequent activation of the UPR pathway (Corazzari et al., 2017). Prolonged ER stress caused by conditions that lead to increased secretory load or the presence of mutated proteins that cannot properly fold in the ER often result in cellular dysfunction and disease (Chadwick and Lajoie, 2019). The UPR and ER stress have also been linked to physiological and premature aging (Hamczyk et al., 2019; Martinez et al., 2017).

Naturally occurring premature aging disorders are powerful model systems to study aging and aging-related pathologies (Gordon et al., 2014; Kubben and Misteli, 2017). One of the most prominent premature aging diseases is Hutchinson-Gilford Progeria Syndrome (HGPS), an extremely rare, fatal genetic condition caused by a *de novo* heterozygous mutation in the *LMNA* gene encoding for the nuclear architectural proteins lamin A and C (De Sandre-Giovannoli et al., 2003; Eriksson et al., 2003; Gordon et al., 2014). The HGPS mutation activates a cryptic splice donor site in *LMNA*, resulting in the expression of progerin, a lamin A isoform that lacks 50 amino acids and permanently retains a C-terminal farnesyl group, leading to its stable association with the inner nuclear membrane (INM) and predominant localization at the nuclear periphery (Gonzalo et al., 2017; Gordon et al., 2014). Progerin acts in a dominant negative fashion and causes a variety of cellular defects including compromised nuclear architecture, heterochromatin maintenance, DNA repair, cell proliferation and differentiation, as well as post-transcriptional reduction of select cellular proteins (Guilbert et al., 2021; Mateos et al., 2013; Pegoraro et al., 2009; Scaffidi and Misteli, 2006; Viteri et al., 2010). Furthermore, progerin has been shown to form static protein aggregates at the periphery of nuclei of patient-derived primary and immortalized fibroblasts (Dahl et al., 2006; Kubben et al., 2016b; McClintock et al., 2006; Vidak et al., 2015). Progerin aggregates are thought to be deleterious due to the entrapment of normal cellular proteins and compromising their function, including the anti-oxidative master regulator NRF2 (Kubben et al., 2016b). Activation of autophagy through rapamycin-mediated mTOR inhibition has been shown to promote the solubilization and clearance of progerin and to alleviate cellular aging defects (Cao et al., 2011), indicating progerin aggregation as one of the drivers of HGPS etiology.

Here we have investigated the cellular chaperone response to the presence of progerin aggregates in HGPS. We find that progerin aggregates at the nuclear periphery promote clustering of the INM transmembrane protein SUN2, which in turn triggers sequestration of chaperones in the ER lumen via the SUN2 luminal domain, leading to induction of ER stress. These observations identify a novel activator of ER stress, a new mechanism of communication between the nucleus and the ER, and they provide insight into the molecular disease mechanisms of HGPS.

## RESULTS

### Progerin alters expression and localization of select molecular chaperones

The premature aging disorder HGPS is caused by the progerin protein, a mutant isoform of the nuclear intermediate filament protein lamin A (De Sandre-Giovannoli et al., 2003; Eriksson et al., 2003; Gordon et al., 2014). Progerin is incorporated into the nuclear lamina and forms insoluble and static protein aggregates with low protein turnover dynamics both at the nuclear periphery and in the nuclear interior (Dahl et al., 2006). We hypothesized that the presence of progerin aggregates leads to activation of a cellular chaperone response.

To assess the effect of progerin on the expression and localization of major molecular chaperones, we performed biochemical and high-throughput imaging analysis to quantitatively determine protein levels and cellular distribution of several chaperones in the well-characterized cellular HGPS model system of hTERT-immortalized skin fibroblasts expressing doxycycline-inducible GFP-progerin and in HGPS patient cells (Fig. S1A) (Kubben et al., 2016a). Upon induction of GFP-progerin for 4 days (Fig. S1A), which is sufficient to induce most major cellular defects associated with HGPS (Pegoraro et al., 2009), the localization of Hsp40, Hsp70 and Hsp90 was unaffected by the presence of progerin (Fig. 1A) and immunoblot analysis did not detect any changes in total protein levels of any of the chaperones at the population level (Fig. S1B, C). However, single cell immunofluorescence analysis using high-throughput spinning-disc confocal microscopy revealed a 1.5 – 1.7-fold increase in average mean levels of Hsc70 and Hsp40 upon GFP-progerin expression (Fig. 1B; P = 0.0077 and P = 0.019, respectively). No significant change in the levels of Hsp90 and Hsp110 was observed (Fig. 1B). Similar observations were made in immortalized HGPS patient skin fibroblasts (Fig. S1B, C), although the increase is less prominent due to the previously described heterogeneity of progerin expression in the patient-derived cell lines (Scaffidi and Misteli, 2008; Vidak et al., 2018). Interestingly, despite unchanged Hsp110 levels, confocal microscopy revealed accumulation of Hsp110 around the nuclei of GFP-progerin expressing cells (Fig. 1C, arrows) and in patient skin fibroblasts with a staining pattern resembling of the ER (Fig. S1F; arrows). Co-staining with the ER-resident chaperone GRP78/BiP confirmed co-localization based on line scan analysis (Fig. 1D; Fig. S1G). Taken together, our data point to changes in the expression levels of the Hsp70/Hsp40 chaperone system, as well as changes in the localization and accumulation of Hsp110 at the ER in the presence of progerin.

**Figure 1.**
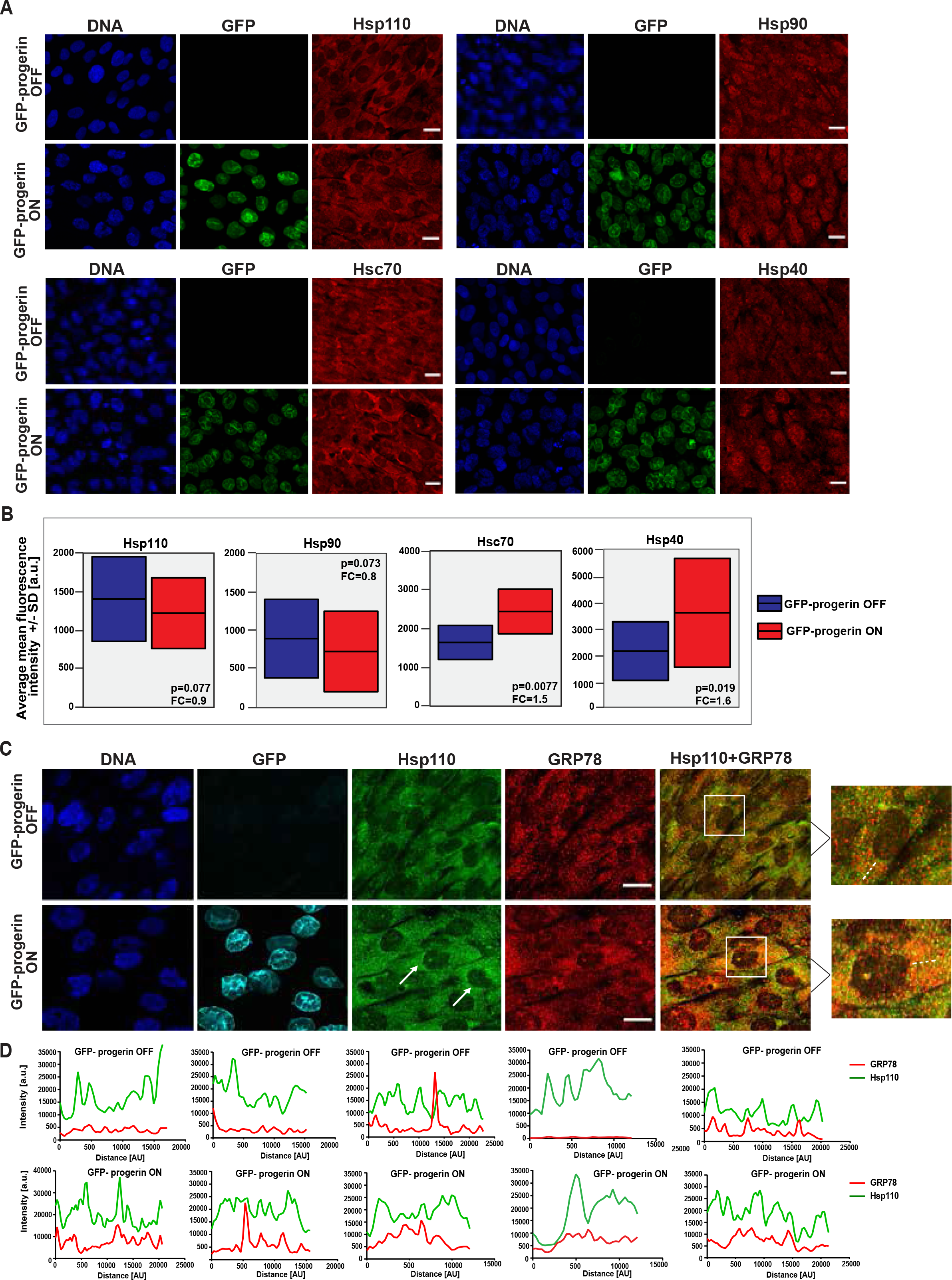
Altered expression and localization of cytosolic chaperones in progerin-expressing cells. **(A)** IF staining of indicated chaperones in PFA fixed GFP-progerin inducible fibroblasts. Scale bar: 10 μm. **(B)** High-throughput IF quantification of chaperone levels in GFP-progerin inducible fibroblasts represented as the average mean fluorescence values ± SD from 3 experiments (number of cells in each experiment = 1300-1500). Statistical differences were analyzed by t-test for the fold change (FC) indicated in the figure. **(C)** IF staining for Hsp110 and GRP78 in PFA fixed GFP-progerin inducible fibroblasts. Arrows indicate Hsp110 accumulation in GFP-progerin expressing cells. GFP panel was false colored aqua marine. Scale bar: 10 μm. **(D)** Line graphs of IF signal intensity along a dotted line based on an overlap in 488 nm and 594 nm channels in 5 randomly selected cells. Progerin induction was for 96 hours (see Materials and Methods).

### Progerin expression induces ER stress *in vitro* and *in vivo*

Disturbances in ER homeostasis result in the accumulation of misfolded proteins within the ER lumen, leading to ER stress (Remondelli and Renna, 2017). These luminal aggregates activate the unfolded protein response (UPR), which alters the expression of numerous genes involved in ER quality control (Lin et al., 2008). In addition, ER stress can result in widespread cytoplasmic protein aggregation and generate proteotoxic stress responses across cellular compartments (Hamdan et al., 2017; Rao and Bredesen, 2004). Given the observed increase in the levels of Hsc70 and Hsp40 chaperones and the accumulation of Hsp110 at the ER in progerin-expressing cells, we sought to determine whether progerin expression causes ER stress by measuring UPR activation *in vitro* and *in vivo*.

Quantitative RT-PCR analysis showed transcriptional activation of several UPR genes, including DDIT3, HSPA5, XBP1 and ERN1/IRE1 upon induction of GFP-progerin (Fig. 2A). The same increase was observed in multiple immortalized and primary HGPS patient fibroblasts (Fig. 2B; Fig. S2A) (Hamczyk et al., 2019). Importantly, induction of these genes was a specific response to the presence of progerin and did not occur upon expression of wild type lamin A (Fig. 2A). These changes represented physiological responses as they were also present in the heart and aorta of homo- and heterozygous progerin transgenic HGPS mice (Varga et al., 2006) and a homozygous knock-in HGPS mouse (Osorio et al., 2011) (Fig. 2C), in line with earlier observations of UPR upregulation in an atherosclerosis-prone HGPS mouse model (Hamczyk et al., 2019). These results were confirmed by single-cell high-throughput imaging which demonstrated a significant increase in the mean protein levels for several ER chaperones including PDI, calnexin, GRP78 and GRP94 in inducible progerin-expressing fibroblasts (Fig. 2D), and a moderate increase in immortalized patient fibroblasts (Fig. 2E). In line with the well established heterogeneity of cellular phenotypes in HGPS cells (Scaffidi and Misteli, 2008; Vidak et al., 2018) single-cell analysis revealed high heterogeneity of expression within the cell population, as well as between biological replicates in HGPS patient cells, with 10-40% of cells showing increase in the expression relative to the wild-type control (Fig. S2B). The same pattern was observed in several primary HGPS patient-derived fibroblasts, with 10-40% cells showing increase in GRP78 and PDI, and up to 80% increase in Calnexin and GRP94 expression levels relative to control (WT 1) (Fig. S2C). Furthermore, immunoblot analysis demonstrated increased phosphorylation of PERK and IRE1α, two major mediators of the UPR pathway, in the presence of progerin (Fig. 2F). We conclude that progerin triggers ER stress pathways.

**Figure 2.**
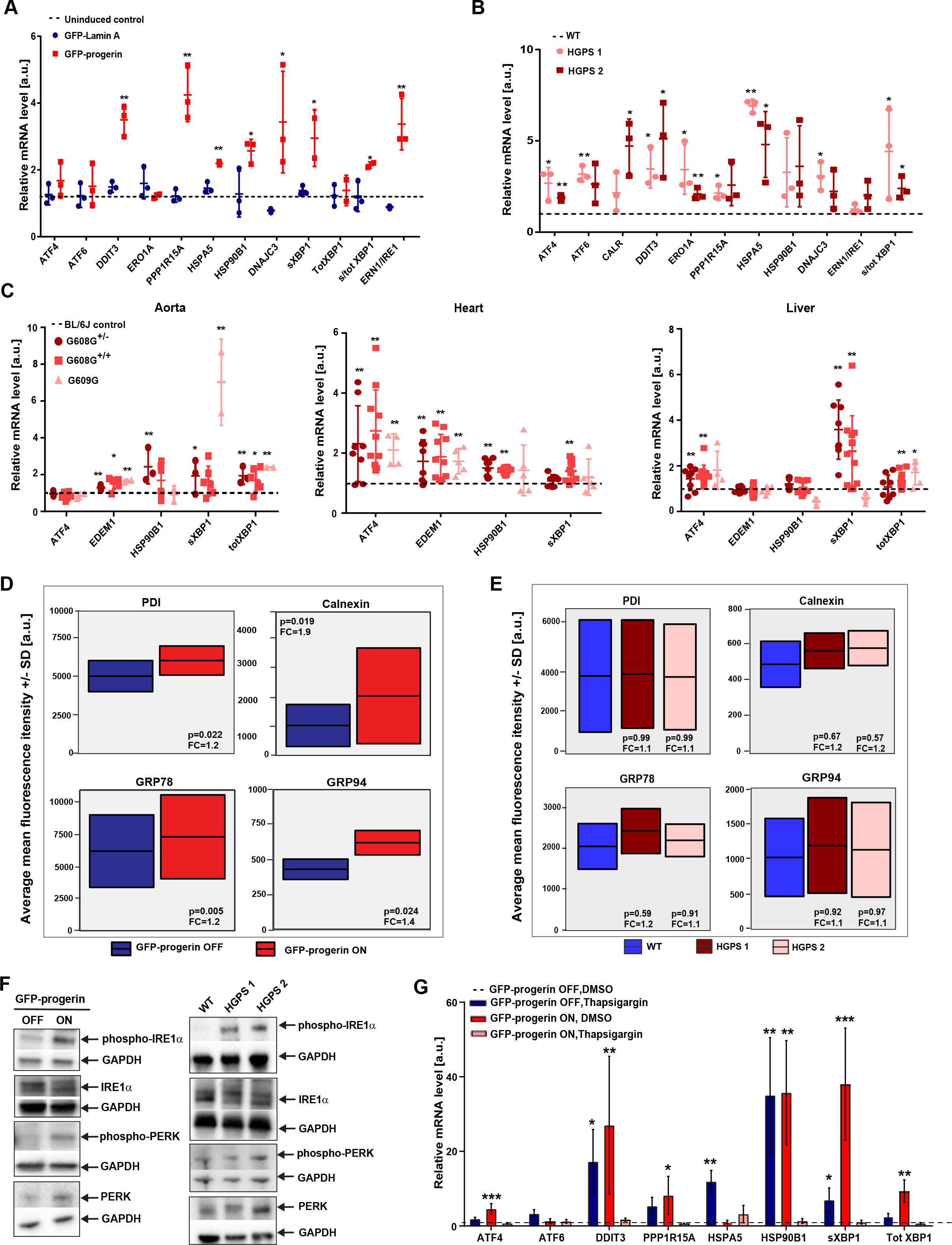
Progerin expression induces ER stress in vitro and in vivo. **(A, B)** Quantification of mRNA levels of UPR genes in GFP-progerin and GFP-lamin A expressing inducible fibroblasts relative to uninduced control **(A)** and in two patient-derived immortalized human HGPS fibroblasts relative to healthy control **(B)** by quantitative qRT-PCR. GAPDH and TBP1 were used for normalization. Values represent the mean of 3 biological replicates ± SD. Statistical differences were analyzed by t-test for GFP-progerin/GFP-lamin A and HGPS/WT. *P<0.05, **P<0.01. Dashed lines represent control value of wild type cells normalized to 1. **(C)** mRNA expression levels of several UPR genes in three different progeria mouse models (G608G^+/-^, G608^+/+^ and G609G^+/+^) relative to C57BL6 control determined by qRT-PCR. GAPDH and TBP1 were used for normalization. Values represent the mean of at least 3 biological replicates ± SD. Dashed lines represent control normalized to 1. Statistical differences were analyzed by t-test between biological replicates. *P<0.05, **P<0.01. **(D)** High throughput IF quantification of ER chaperone expression levels in GFP-progerin inducible fibroblasts represented as the average mean fluorescence values ± SD from 3 experiments (number of cells per experiment >300). Statistical differences were analyzed by t-test for the fold change (FC) indicated in the figure. **(E)** Single cell high throughput IF quantification of ER chaperone expression levels in immortalized human HGPS fibroblasts (number of cells per experiment >300) represented as the distribution of average mean fluorescence intensity values from 3 experiments ± SD (See Materials and Methods). Statistical differences were analyzed by one-way ANOVA followed by the Dunnett’s test using the WT cell line as the negative control**. (F)** Immunoblot analysis of phospho-IRE1α, total IRE1α, phospho-PERK and total PERK levels in total cell lysates of GFP-progerin inducible and patient-derived fibroblasts. GAPDH and γ-tubulin served as a loading control. **(G)** qRT-PCR analysis of mRNA levels of several UPR genes in GFP-progerin inducible fibroblasts treated with Thapsigargin or DMSO (vehicle control). Values for GFP-progerin OFF/Thapsigargin and GFP-progerin ON/DMSO were calculated relative to GFP-progerin OFF/DMSO control, whereas GFP-progerin ON/Thapsigargin is shown relative to GFP-progerin ON/DMSO. GAPDH and TBP1 were used for normalization. Values represent means of 3 biological replicates ± SEM. Statistical differences were analyzed by t-test. *P<0.05, **P<0.01.

### Chronic activation of ER stress in HGPS cells

Elevated levels of ER chaperones and transcriptional activation of the UPR genes are typically associated with a heightened ER stress response (Halperin et al., 2014). To assess whether the progerin-mediated increase in key ER stress response factors affects the ability of cells to respond to stress, we challenged progerin-expressing cells with the ER stressors Thapsigargin or Tunicamycin. Remarkably, while control cells were able to significantly upregulate UPR genes progerin-expressing cells failed to respond to additional ER stress above their elevated baseline level of UPR activity (Fig. 2G; Fig. S2D). These results suggest that progerin chronically activates ER stress response pathways to a level that prevents further activation upon additional stressors.

Previous studies have shown that tauroursodeoxycholic acid (TUDCA), an inhibitor of ER stress, can significantly relieve ER stress and reduce ER stress-mediated cell death (Choi et al., 2016a; Choi et al., 2016b; Uppala et al., 2017; Zhang et al., 2018). Treatment of GFP-progerin inducible cells with 25 μM TUDCA for 48h significantly increased cell number in comparison to vehicle treated control (Fig. S2B) but had no effect on progerin levels (Fig. S2E) nor total GRP78 levels (Fig. S2G), consistent with a role of ER stress in progerin-mediated cell death, one of the major HGPS cellular phenotypes (Vidak et al., 2015; Zhang et al., 2014). Together with recent findings demonstrating beneficial effects of TUDCA on vascular phenotype and lifespan of atherosclerosis-prone progeria mice (Hamczyk et al., 2019), our data suggest a contribution of ER stress to the development of the HGPS phenotypes.

### Progerin aggregates sequester ER chaperones at the nuclear periphery

To investigate the mechanism by which the nuclear protein progerin induces ER stress, we probed the distribution of several ER chaperones in progerin-expressing cells using super-resolution structured illumination microscopy (SIM) (Fig. 3). We find accumulation of GRP78/BiP and PDI in ER regions immediately juxtaposed to the nuclear periphery, mirroring the distribution of progerin (Fig. 3A, B). Quantification of single-cell high-throughput imaging of GRP78/BiP distribution shows significant recruitment to the nuclear periphery in both GFP-progerin expressing and patient-derived fibroblasts, as assessed by measuring the mean fluorescence intensity of a band of 5-35% distance from the nucleus, where 0% is adjacent to DAPI and 100% represents the cell border (Fig. 3C, D; see Materials and Methods). Expression of wild-type lamin A did not induce any changes in GRP78/BiP localization or UPR activity (Fig. S3 A-C). Co-immunoprecipitation of GRP78/BiP with progerin, but not with wild type lamin A (Fig. 3E, Fig. S3D), confirmed physical association, most likely indirectly (see below), of progerin with the resident ER chaperone GRP78/BiP.

To determine whether the accumulation of GRP78/BiP at the nuclear periphery was a direct consequence of the presence of progerin aggregates at the nuclear envelope or was an indirect, non-specific stress response, we expressed the non-farnesylated C661S mutant of progerin, which accumulates in the nucleoplasm but not at the nuclear periphery (Fig. 3F) (Kubben et al., 2016a). Progerin-C661S, in contrast to progerin, does not lead to significant GRP78/BiP sequestration (Fig. 3G) nor to the induction of UPR signaling or increased levels of ER chaperones (Fig. 3H, I). In addition, co-immunoprecipitation of GRP78/BiP with Progerin-C661S showed no physical association between non-farnesylated progerin and GRP78/BiP (Fig. 3J; Fig. S3D). These results demonstrate that the presence of progerin aggregates at the nuclear periphery triggers GRP78/BiP sequestration to the ER regions juxtaposed to the nuclear envelope and to the induction of ER stress.

**Figure 3.**
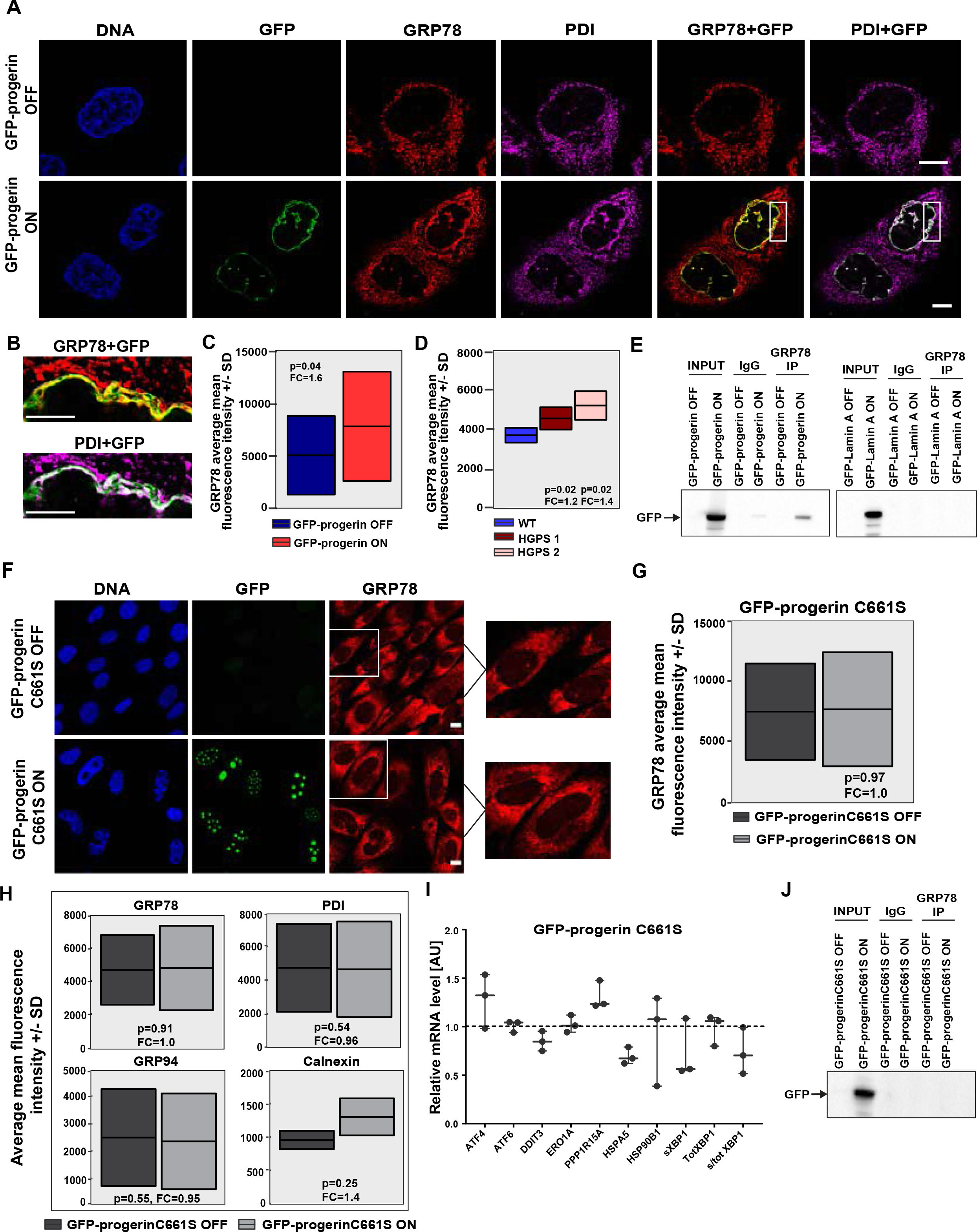
ER chaperones accumulate around the nuclear periphery of progerin-expressing cells. **(A)** Representative structured illumination microscopy (SIM) images of GRP78 and PDI staining in GFP-progerin inducible fibroblasts fixed with PFA. Scale bar: 10μm. **(B)** Enlarged view of the nuclear periphery in GFP-progerin inducible fibroblasts from panel A. Scale bar: 5μm. **(C, D)** High throughput IF quantification of GRP78 recruitment to the nuclear periphery in GFP-progerin inducible fibroblasts **(C)** and in patient-derived HGPS fibroblasts **(D)** assessed by measuring the mean fluorescence intensity of a band of 5-35% distance from the DAPI-based nuclear region of interest, where 0% is adjacent to DAPI and 100% represents the cell boundary (number of cells for each experiment >15001500). Data is represented as the average mean fluorescence values ± SD from 3 experiments (See Materials and Methods). Statistical differences were analyzed by t-test for the fold change (FC) indicated in the figure. **(E)** Western blot analysis of interaction between immunoprecipitated GRP78 with GFP-lamin A or GFP-progerin in inducible fibroblasts. **(F)** GRP78 IF staining in PFA fixed non-farnesylated GFP-progerin C661S inducible fibroblasts (see Materials and Method). Scale bar: 10μm. **(G)** High throughput IF quantification of GRP78 recruitment to the nuclear periphery in non-farnesylated GFP-progerin inducible fibroblasts (number of cells for each experiment >1500) shown as the average mean fluorescence values ± SD from 3 experiments. Statistical differences were analyzed by t-test for the fold change (FC) indicated in the figure. **(H)** High-throughput IF quantification of ER chaperone expression levels in non-farnesylated GFP-progerin inducible fibroblasts (number of cells for each experiment >1500) represented as the average mean fluorescence values ± SD from 3 experiments. Statistical differences were analyzed by t-test for the fold change (FC) indicated in the figure. **(I)** qRT-PCR analysis of mRNA levels of several UPR genes in non-farnesylated GFP-progerin inducible fibroblasts relative to uninduced control. GAPDH and TBP1 were used for normalization. Dashed lines represent control normalized to 1. Values represent means of 3 biological replicates ± SD. Statistical differences were analyzed by t-test. *P<0.05, **P<0.01. **(J)** Western blot analysis of interaction between immunoprecipitated GRP78 with GFP-progerinC661S in inducible fibroblasts.

### The inner nuclear membrane protein SUN2 mediates the ER stress phenotype triggered by nuclear aggregation

Progerin is a nucleoplasmic protein tightly associated with the nuclear envelope (Dahl et al., 2006), yet it appears to trigger the UPR in the ER lumen and can biochemically be pulled down with the ER-luminal chaperone GRP78/BiP (see Fig. 3). However, direct interaction between progerin and ER chaperones is unlikely since progerin does not extend into the perinuclear space, pointing to the existence of adaptor proteins that sense and transmit the presence of nucleoplasmic progerin aggregates to the ER lumen. Because the ER lumen is contiguous with the perinuclear space between the INM and outer nuclear membrane (ONM), we hypothesized that the INM proteins SUN1 and SUN2 may mediate the nuclear-triggered ER stress response by transmitting the activating signal across the INM into the ER lumen (Starr and Fridolfsson, 2010; Wang et al., 2012). SUN1 and SUN2 are transmembrane proteins whose N-terminus interacts with the nuclear lamin proteins, including progerin, on the nucleoplasmic face of the nuclear envelope, whereas their highly conserved luminal C-terminus extends into the perinuclear space where it interacts with nesprin transmembrane proteins located in the ONM (Haque et al., 2010). As part of the Linker of Nucleoskeleton and Cytoskeleton complex (LINC), which spans the lumen of the NE, SUN proteins act as signal transmitters between the nucleus and the cytoplasm, in particular of mechanical forces (Hieda, 2019). Interestingly, SUN1/2 have been linked to progeric and dystrophic laminopathies (Chen et al., 2012).

In line with a role for SUN proteins in sensing of progerin aggregates and transmitting their presence to the ER lumen, single-cell high-throughput imaging revealed that both SUN1 and SUN2 levels were increased upon GFP-progerin expression (Fig. S4A), as well as in various immortalized and primary patient-derived fibroblast cell lines (Fig. S4B, C, D). Furthermore, SIM revealed accumulation and co-localization of progerin, SUN1 and SUN2 with GRP78/BiP or PDI at the nuclear periphery in GFP-progerin expressing cells (Fig. 4A, B; Fig. S4E, F and G), as well as the local enrichment of SUN2, GRP78 and PDI in the presence of progerin aggregates (Fig. 4C). Similar co-localization was observed in highly lobulated nuclei of patient-derived fibroblasts as assessed by confocal microscopy, line scan analysis and measuring local enrichment of SUN2, GRP78 and PDI at the nuclear periphery (Fig. S4H, I and J). Inhibiting the ability of inner nuclear membrane SUN proteins to interact with KASH proteins in the outer nuclear membrane does not impair the ability of progerin to activate ER stress (Fig. S4K), as shown by the expression of a dominant negative KASH domain (DN-KASH), which disrupts SUN2 interactions (Lombardi et al., 2011).

**Figure 4.**
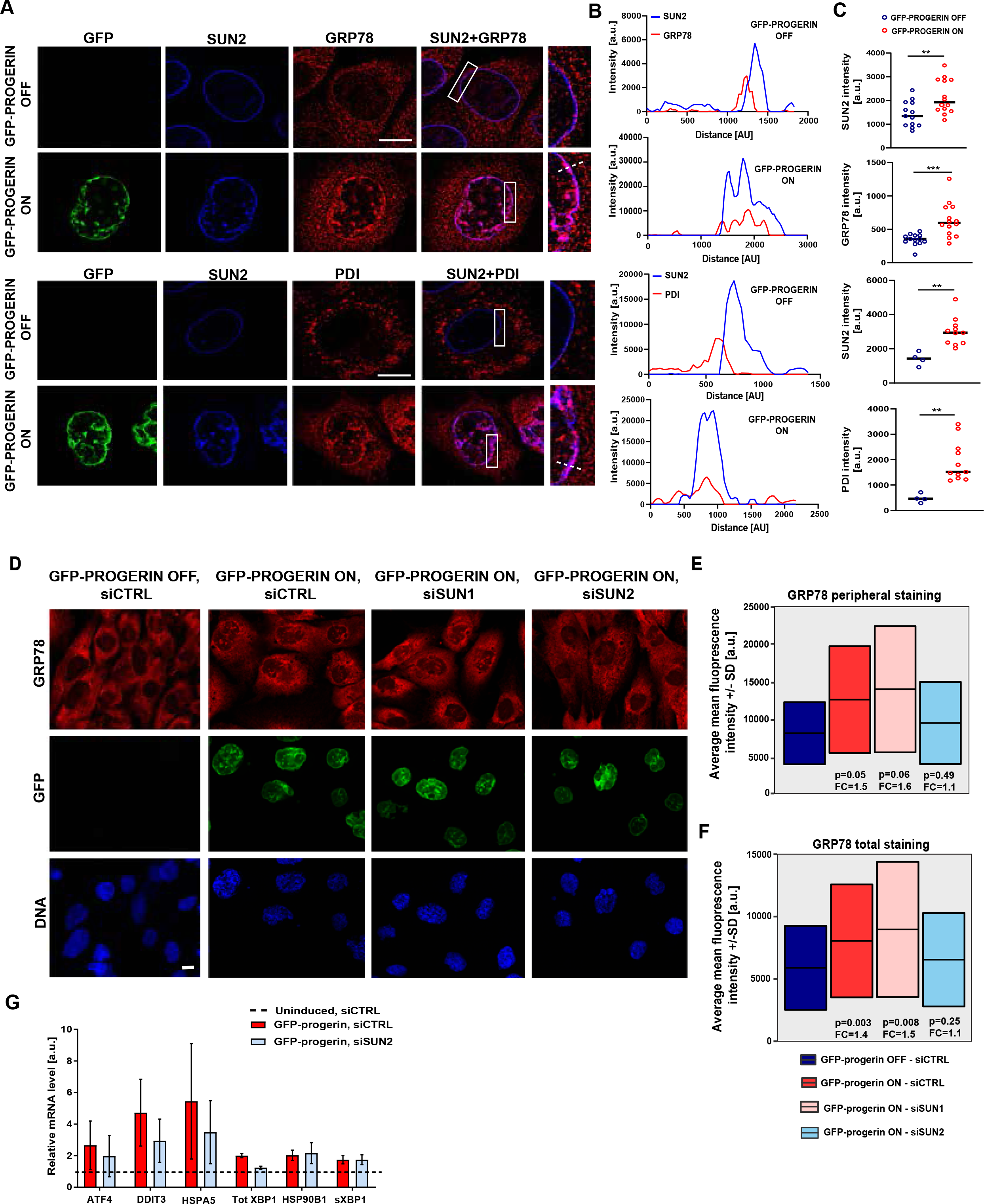
SUN2 contributes to the ER stress phenotype in progerin-expressing cells. **(A)** Representative SIM images of GRP78, PDI and SUN2 staining in GFP-progerin inducible fibroblasts fixed with PFA. Scale bar: 10μm. **(B)** Line graphs of IF signal intensity across the dotted lines in panel A based on an overlap in 594nm and 647nm channels. **(C)** Scatter plot of fluorescence intensities of the peripheral SUN2, GRP78 and PDI localization (n=4-15). Line represents the median. Statistical differences were analyzed by t-test. **(D)** Confocal images of GRP78 IF staining in siSUN1 and siSUN2 treated GFP-progerin inducible fibroblasts and their respective controls. Scale bar: 10μm. (**E, F**) High throughput IF quantification of GRP78 recruitment to the nuclear periphery **(E)** and total GRP78 levels **(F)** in siSUN1 and siSUN2 treated GFP-progerin inducible fibroblasts and their respective controls (number of cells for each experiment = 200-400) represented as the average mean fluorescence values ± SD from 3 experiments. Statistical differences were analyzed by t-test for the fold change (FC) indicated in the figure. **(G)** qRT-PCR analysis of mRNA levels of several UPR genes in siSUN2 and control treated GFP-progerin inducible fibroblasts relative to uninduced control. GAPDH and TBP1 were used for normalization. Dashed lines represent control normalized to 1. Values represent means of 3 biological replicates ± SD. Statistical differences were analyzed by t-test. *P<0.05, **P<0.01.

To determine whether SUN1 and SUN2 indeed contribute to the progerin-mediated accumulation of ER proteins at the nuclear periphery, we performed siRNA knockdown of each SUN protein and measured the recruitment of GRP78/BiP to the nuclear periphery upon induction of GFP-progerin expression (Fig. 4D, E; Fig. S5A, B). While loss of SUN1 had no effect on GRP78/BiP accumulation at the nuclear periphery, knockdown of SUN2 prevented GRP78/BiP accumulation and restored normal GRP78/BiP localization even when progerin was induced (Fig. 4D, E). In addition, SUN2, but not SUN1, downregulation decreased total GRP78/BiP levels (Fig. 4F) and prevented the progerin-mediated transcriptional upregulation of UPR genes (Fig. 4G). siRNA knockdown of each SUN protein had no effect on the total GRP78/BiP levels or GRP78/BiP accumulation at the nuclear periphery in uninduced control (Fig. S5C). We conclude that the progerin-interactor SUN2 contributes to the ER stress phenotype observed when progerin aggregates are present in the nucleoplasm.

### SUN2 clustering at the nuclear periphery activates the UPR response through the recruitment of GRP78/BiP

To uncover how protein aggregates at the nucleoplasmic face of the nuclear envelope can cause a change in the localization of ER chaperones and activate ER stress pathways via SUN2, we analyzed the expression and the localization of SUN2 in more detail. Stimulated emission depleted (STED) super-resolution microscopy reveled that SUN2 proteins are not uniformly localized at the nuclear periphery but accumulate in cluster-like structures in the INM in the presence of progerin but not of wild type lamin A (Fig. 5A, arrows). Interestingly, SUN2 clusters coincide with the highest levels of GRP78/BiP accumulation (Fig. 5A, arrows). The accumulation of SUN2 was not merely a reflection of higher local levels of progerin, since the ratio of SUN2 relative to progerin increased compared to wild type lamin A, pointing to local enrichment of SUN2 in the presence of progerin aggregates (Fig. 5B). Similarly, GRP78/BiP was enriched at sites of SUN2 clusters induced by progerin aggregation (Fig. S6A), indicating recruitment of GRP78/BiP in the INM lumen to the newly formed SUN2 clusters.

**Figure 5.**
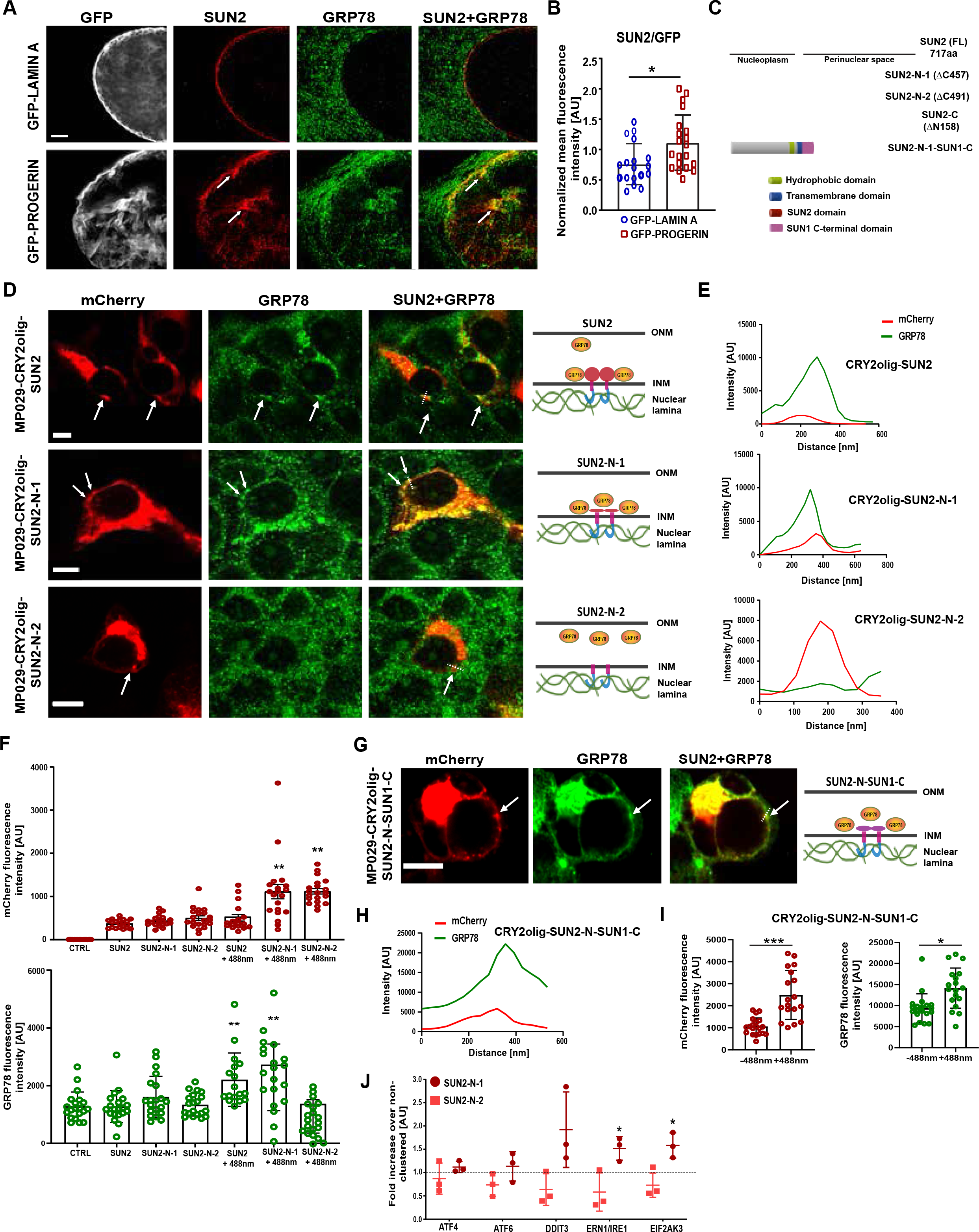
SUN2 clustering causes changes in the localization of ER chaperones. **(A)** Representative stimulated emission depletion microscopy (STED) images of GRP78 and SUN2 staining in GFP-lamin A and GFP-progerin inducible fibroblasts. Arrows indicate SUN2 and GRP78 clusters. Scale bar: 2μm. **(B)** SUN2 mean fluorescence intensity normalized to the GFP signal in GFP-lamin A (n=18) and GFP-progerin (n=20) inducible fibroblasts. The measurements were done on maximum projection STED images by measuring the whole ROI. Statistical differences were analyzed by t-test. *P<0.05. **(C)** Schematic representation of SUN2 and its deletion constructs. The conserved SUN domain is depicted in red, the transmembrane segment in blue, additional hydrophobic stretches in green and C-terminal SUN1 fragment in purple. **(D)** Representative confocal images of MP029-CRY2olig-mCherry SUN2 variants driven by the ubiquitin-promoter driven for low level expression of SUN2 and stained for GRP78 in HEK293FT cells fixed with PFA after cluster induction. Scale bar: 10μm. Arrows indicate SUN2 and GRP78 clusters. Drawings on the right depict localization of SUN2 variants in the INM and their level of GRP78 recruitment. **(E)** Line graphs of IF signal intensity across the clusters (dashed line) based on an overlap in 488 nm and 594 nm channels in panel D. **(F)** Mean fluorescence intensities of the mCherry and GRP78 in non-clustered and clustered cells (+488nm) transfected with various MP029-CRY2mCherry constructs (n=20). Statistical differences were analyzed by t-test relative to no plasmid control. **P<0.005. **(G)** Representative confocal images of the MP029-CRY2olig-mCherry SUN2-N variant fused to SUN1 C-terminal fragment (230aa) expressed in HEK293FT cells after cluster induction. Cells were fixed with PFA and stained with GRP78 antibody. Scale bar: 10 μm. Arrows indicate SUN2-N-SUN1-C and GRP78 clusters. Drawings on the right depict localization of SUN2-N-SUN1-C variant in the INM and its level of GRP78 recruitment. **(H)** Line graphs of IF signal intensity across the clusters (dashed line) based on an overlap in 488nm and 594 nm channels in panel G. **(I)** Mean fluorescence intensities of the mCherry and GRP78 in non-clustered and clustered cells (+488nm) transfected with MP029-CRY2olig-mCherry SUN2-N-SUN1-C construct (n=20). Statistical differences were analyzed by t-test relative to non-clustered control. *P<0.05, **P<0.005. **(J)** qRT-PCR analysis of mRNA levels of several UPR genes in HEK293FT cells transfected with MP029-CRY2olig-mCherry SUN2-N variants. Data is shown as fold increase of cluster-induced over non-induced samples. Constructs were under control of a ubiquitin promoter for low level expression. GAPDH and TBP1 were used for normalization. Dashed lines represent non-transfected control used to normalize the data (set to 1). Values represent means of 3 biological replicates ± SD. Statistical differences were analyzed by t-test. *P<0.05, **P<0.01.

The requirement for SUN2 in activation of the luminal ER stress response combined with the observed local clustering of SUN2 and the accumulation of GRP78/BiP at SUN2 clusters suggests a mechanistic model in which SUN2 senses the density of lamin A/progerin on the nucleoplasmic side of the INM and transmits this information to the ER lumen via its clustering (Fig. 5A; Fig. S6B). In this model SUN2 detects the presence of progerin aggregates via its known interaction of its nucleoplasmic N-terminus with progerin, while SUN2 clustering is sensed in the ER lumen by interaction of GRP78/BiP with the clustered SUN2 C-terminal domain (Fig. S6B). This model predicts that forced clustering of SUN2 in the absence of progerin aggregates should be sufficient to recruit GRP78/BiP to SUN2 clusters in the ER and that this accumulation should be dependent on the C-terminal domain of SUN2 (Fig. S6B). To test these predictions, we developed an optimized optogenetic clustering approach based on the *Arabidopsis thaliana* CRY2-derived optogenetic module CRY2olig, which induces rapid and robust protein oligomerization upon exposure to blue light (Taslimi et al., 2014). Several SUN2 variants (Fig. 5C) were fused to CRY2olig-mCherry and expressed under the control of either a strong CMV promoter or a ubiquitin promoter for moderate to low expression. As expected based on prior observations on SUN2 localization upon exogenous expression (Ungricht et al., 2015), high level expression of wild-type SUN2 or mutants that contained the transmembrane domain and either the complete (SUN2-C) or partial luminal domain (SUN2-N-1) resulted in the formation of cluster-like structures even in the absence of light induction (Fig. S6C). Interestingly, line scan analysis by confocal microscopy revealed accumulation of GRP78/BiP at these SUN2 clusters, with the most prominent accumulation at clusters of SUN2-C containing the full luminal domain (Fig. S6C, D, E). No such accumulation was observed upon expression of SUN2 at low levels (Fig. 5F; Fig. S6F, G). Overexpression of any of the SUN2 variants did not impact cell proliferation and did not induce any cellular progeric phenotypes (Fig. S6H). These data suggest that high level expression of SUN2 induces its clustering and promotes GRP78/BiP sequestration to the nuclear periphery via the luminal portion of the protein.

To directly demonstrate that clustering of SUN2 triggers accumulation of ER chaperones, we induced SUN2 clustering in a controlled fashion upon low level expression of CRY2olig-SUN2 driven by the ubiquitin promoter by optogenetics using ten 200ms light pulses over 90min (see Materials and Methods). As previously described, live cell imaging revealed rapid clustering of CRY2olig-mCherry (Fig. S7A, panel 1) (Taslimi et al., 2014). Consistent with slow lateral diffusion dynamics of SUN proteins in the INM (Liu et al., 2007; Ungricht et al., 2015), the full length SUN2 started to re-organize over 30 min and reached full clustering at 90 min (Fig. S7A, panel 2). In line with the ability of clustered SUN2 to trigger ER stress responses, GRP78/BiP accumulated strongly at light-induced SUN2 clusters at the nuclear periphery in the absence of progerin (Fig. 5D, E; top panel, Fig. 5F), recapitulating the phenotype observed in progerin-expressing cells. To demonstrate that it is the clustering of the luminal portion of SUN2 that promotes GRP78/BiP sequestration, we optogenetically induced clustering of SUN2-N-1 and SUN2-N-2 variants, which lack part or all of the luminal domain, respectively (Fig. 5C). These mutants clustered more readily than full-length SUN2 (Fig. S7A, panels 3 and 4), most likely due to significantly smaller sizes of these proteins (260 aa and 226 aa, respectively, in comparison to 717 aa for the full length SUN2). Fluorescence Recovery After Photobleaching (FRAP) analysis shows very little recovery of induced clusters with the recovery rate of 0.086 +/-0.0075 s-1 (Fig. S7B). Confocal microscopy revealed that clusters of SUN2-N-1, which contains a short portion of the luminal domain, still recruit GRP78/BiP to the nuclear periphery (Fig. 5D, middle panel, see arrows; Fig. 5E, F). In contrast, induced clusters of SUN2-N-2, which completely lacks the luminal domain, do not recruit GRP78/BiP (Fig. 5D, lower panel; Fig. 5E, F). These data demonstrate that the entire luminal portion of the SUN2 protein is necessary and part of the luminal domain are sufficient for the recruitment of the GRP78/BiP to the nuclear periphery. Interestingly, clustering of a construct consisting of the nucleoplasmic and transmembrane domains of SUN2 fused to the luminal domain of SUN1 (Fig. S7A, panel 5) still efficiently recruits GRP78 to the induced clusters (Fig. 5G, H, I), suggesting that GRP78/BiP recognition of clusters is sequence-independent.

To finally show that the peripheral recruitment of the GRP78/BiP leads to the activation of the UPR response, we enriched mCherry positive cells by fluorescence-activated cell sorting (FACS) (Fig. S7B and C) and performed qRT-PCR analysis of UPR genes upon optogenetic clustering of SUN2 variants (see Materials and Methods). Optogenetically-induced clustering was confirmed by immunofluorescence analysis prior to qRT-PCR analysis (Fig. S7C and D). qRT-PCR analysis showed transcriptional activation of several UPR genes, including DDIT3, ERN1/IRE1 and EIF2AK3 upon induction of SUN2-N-1 clusters (1.5 – 1.9-fold increase), whereas clustering of the SUN2-N-2 variant completely lacking the C-terminal luminal domain did not further increase expression of UPR genes (Fig. 5F, Fig. S7E). These data demonstrate that clusters of SUN2 protein in the INM lead to the activation of the UPR response.

## DISCUSSION

We identify here, in the context of the premature aging disorder HGPS, a novel mechanism of activation of ER stress response pathways. We find that the INM transmembrane protein SUN2 senses the aggregation of the disease-causing progerin protein at the nucleoplasmic face of the nuclear envelope and transmits this information to the ER lumen to trigger the UPR. These findings reveal an unanticipated pathway of communication between the nucleus and the ER and they provide insights into the molecular disease mechanisms of HGPS.

ER stress response pathways, including the UPR, are triggered by the accumulation of misfolded and unfolded proteins in the ER lumen (Preissler and Ron, 2019). The UPR is activated when the major ER-resident chaperone GRP78/BiP dissociates from the major UPR transducers IRE1α, PERK and ATF6 and binds to unfolded proteins in the ER lumen (Gardner et al., 2013). The UPR can be triggered by numerous stimuli including redox disruption, hypoxia, aberrant Ca^2+^ regulation, energy balance disruption, viral infections, overproduction of secreted/transmembrane proteins, secretion of folding-incompetent mutated proteins, as well as a number of xenobiotics (Lafleur et al., 2013). Our observations expand these mechanisms of UPR activation by demonstrating that ER stress pathways can also be triggered by nucleoplasmic protein aggregates at the periphery of the nucleus. We show that the transmembrane INM protein SUN2 senses progerin aggregates juxtaposed to the nuclear envelope via interactions of its N-terminus with progerin, leading to SUN2 clustering in the INM. Analogous to typical UPR activation, clustering of SUN2 is then sensed by GRP78/BiP in the perinuclear space, which is contiguous with the ER lumen, via recognition of the C-terminal luminal regions of SUN2, thus triggering the UPR pathway. This signaling axis represents a novel mechanism of ER stress activation induced by a nuclear trigger. Whether other nuclear proteins can trigger similar ER stress by the same or related mechanisms remains to be determined.

Several lines of evidence support the conclusion that clustering of the luminal portion of SUN2 induces recruitment of the GRP78/BiP to SUN2 clusters and activation of the UPR response. First, artificial clustering by optogenetic methods of the full length SUN2 or a SUN2 N-terminus variant that still contains a portion of the C-terminus is sufficient to promote GRP78/BiP sequestration and transcriptional activation of several UPR genes. Second, overexpression of the SUN2 C-terminus which consists of the luminal and transmembrane domain of SUN2 induces pronounced cluster-like structures which very efficiently recruit GRP78/BiP, similar to the high-density aggregates observed in progerin expressing cells. Third, optogenetic clustering of a SUN2 variant which completely lacks the C-terminal luminal domain does not recruit GRP78/BiP and does not induce transcriptional activation of UPR genes, in contrast to full length SUN2 or SUN2-N-1 induced clusters. The observation that SUN2-N-1 variant containing only a small portion of the luminal domain (34aa) still efficiently recruits GRP78/BiP demonstrates that GRP78/BiP recognition of clusters is a sequence-independent effect, as confirmed by clustering the construct consisting of nucleoplasmic and transmembrane domains of SUN2 fused to the luminal domain of SUN1.

Our results document a communication pathway between the nuclear interior and the ER lumen and as such establish a new mechanism of inter-compartment communication. We identify the inner nuclear membrane protein SUN2 as a mediator of this communication. SUN2, together with SUN1, is a member of the SUN (Sad1p, Unc-84) domain family of proteins (Malone et al., 1999) which share a conserved SUN domain at their luminal C-terminus (Tzur et al., 2006), at least one transmembrane domain and a less characterized N-terminus which interacts with the nuclear lamin A protein on the nucleoplasmic face of the INM (Haque et al., 2010). SUN proteins interact in the lumen of the nuclear envelope with the outer nuclear membrane KASH (Klarsicht, Anc-1, Syne-1 homology) domain proteins to form the core of the LINC complex involved in connecting the nuclear lamina and the cytoskeleton (Kim et al., 2015). As such, the LINC complex plays an important role in mechanotransduction, nuclear migration, cell polarity, cytoskeletal organization, meiotic chromosome movement and protein recruitment to the ONM (Kim et al., 2015). Our findings expand the mode of action of SUN proteins by demonstrating that SUN2 does not only function via interaction with the KASH-domains in nesprins. Whether the interaction of SUN2 with GRP78/BiP in the presence of progerin reflects an amplification of more sporadically occurring binding events in a non-disease state or whether the interaction is specifically triggered under pathological conditions by the presence of progerin remains to be determined.

The ability of SUN2 to sense the aggregation of progerin is based on its intrinsic propensity to interact with nuclear lamins as part of the LINC complex (Haque et al., 2010). In HGPS, the disease-causing progerin protein becomes integrated into the nuclear lamina and accumulates at the nuclear periphery leading to thickening of the lamina and formation of static protein aggregates (Dahl et al., 2006). The presence of these nucleoplasmic, membrane proximal aggregates results in abnormal clustering of SUN2 and activation of ER stress responses. In addition to the interaction of the SUN2 nuclear domain with lamins, clustering is likely also facilitated by the intrinsically low mobility of SUN proteins in the INM (Liu et al., 2007). Our finding that SUN2 acts as a sensor of nuclear protein aggregates, transmits the signal to the ER, and activates stress response pathways to restore homeostasis describes a new function for a protein in the LINC complex.

Sustained ER stress and a faulty UPR response have been linked to physiological aging, as well as to many aging-associated diseases (Brown and Naidoo, 2012). The naturally occurring premature aging disorder HGPS mimics several aspects of normal aging, including increased DNA damage, epigenetic changes and the emergence of vascular defects (Kubben and Misteli, 2017). Our work now extends these parallels by describing activation of the ER stress response in progerin-expressing cells, in HGPS patient cells as well as in several progeria mouse models. These findings complement the recent observation of transcriptional misregulation of ER stress and UPR components in two mouse models of HGPS with ubiquitous or vascular smooth muscle cell-specific progerin expression (Hamczyk et al., 2019) pointing to a significant role of ER stress pathways in HGPS. Interestingly, chronic activation of UPR as well as the inability to activate it upon challenge as found in HGPS cells has previously been associated with normal aging (Taylor and Hetz, 2020). In adition, previous studies have shown that nuclear irregularities in HGPS fibroblasts correlate with SUN1 overexpression and knocking down of SUN1 alleviates some of the HGPS nuclear defects (Chen et al., 2012). We show here that SUN2 overexpression in wild type cells does not recapitulate HGPS cellular phenotypes, but rather contributes to the disease development by chronic activation of the ER stress response, ultimately leading to cell death. These findings demonstrate distinct functions for SUN1 and SUN2 in HGPS, which is in line with prior findings demonstrating both redundant and separate functions for SUN isoforms depending on their context (Lei et al., 2009; Thakar et al., 2017).

Our finding of UPR activation in HGPS may also have therapeutic implications. Treatment with the chemical chaperone TUDCA has previously been reported to efficiently alleviate ER stress in a number of HGPS-relevant physiological systems including models of aortic valve calcification, hypertrophy and diabetes (Cai et al., 2013; Ozcan et al., 2006; Rani et al., 2017). Furthermore, treatment with TUDCA ameliorates vascular pathology and prolongs lifespan in atherosclerosis-prone HGPS mouse models (Hamczyk et al., 2019). These findings uncover an unexplored aspect of this premature aging disease identifying ER stress and the UPR as drivers of cardiovascular and cellular phenotypes in HGPS and they suggest that alleviation of ER stress may offer a novel therapeutic strategy for HGPS.

Taken together, our observations delineate a novel cellular stress sensor pathway linking the nucleus to the ER and they suggest that improper protein sequestration at the nuclear periphery may be responsible for prominent defects associated with premature and possibly physiological aging.

## MATERIALS AND METHODS

### Cell culture and treatments

hTERT-immortalized GFP-lamin A, GFP-progerin, and GFP-progerin C661S doxycycline inducible dermal fibroblast cell lines were maintained and induced (96 hr) as described (Kubben et al., 2016a). Immortalized patient-derived HGPS (72T=HGPS 1; 97T=HGPS 2) and age-matched wild-type control (CRL-1474) cell lines were cultured as described (Fernandez et al., 2014; Hahn et al., 1999; Scaffidi and Misteli, 2011). Experiments in immortalized cells were performed within the first 15 passages after transformation. Cells were grown in MEM containing 15% fetal bovine serum (FBS), 2 mM l-glutamine, 100 U ml^− 1^ penicillin, 100 μgml^− 1^ streptomycin, 1 mM sodium pyruvate and 0.1 mM non-essential amino acids at 37 °C in 5% CO_2_. Primary human dermal fibroblast cell lines were obtained from the Progeria Research Foundation (PRF) and National Institute of Aging (NIA) Cell Repository distributed by the Coriell Institute. Four of the cell lines were from healthy donors and five were from patients that have the classic mutation in *LMNA* Exon 11, heterozygous c.1824C > T (p.Gly608Gly). The list of the cell lines used together with the source and the age at donation are listed in Table S1. For all experiments, cells were passage matched between P10-20. Cells were grown in high glucose DMEM containing 15% fetal bovine serum (FBS), 1% GlutaMAX (ThermoFisher #35050-061) and 1% Penicillin-Streptomycin (ThermoFisher #15140-122) at 37 °C in 5% CO_2_. U-2 OS cells (ATCC® HTB-96™) and HEK293FT cells (Thermo Fisher Scientific, R70007) were grown in DMEM containing 10% fetal bovine serum (FBS), 2 mM l-glutamine, 100 U ml^− 1^ penicillin and 100 μgml^− 1^ streptomycin, at 37 °C in 5% CO_2_. All cell lines were mycoplasma negative as shown by routine testing (EZ-PCR™ Mycoplasma Detection Kit, Biological Industries).

Cell treatments included 1μM Thapsigargin (Sigma, T9033), 1μM Tunicamycin (Sigma, T7765), 25μM TUDCA (Milipore, #580549) and their respective vehicle controls. Thapsigargin was dissolved in DMSO, Tunicamycin in EtOH and TUDCA in sterile cell-culture grade H2O. Cells were treated with Thapsigargin and Tunicamycin for 2h and with TUDCA for 48h at 37 °C.

### Plasmid construction and expression

To create CRY2oligo-mCherry-SUN2 (human), a *BSRGI* site was added N-terminal to the SUN2 sequence in the MYC-BioID2-SUN2 construct, a kind gift from Dr. Kyle Roux (Sanford School of Medicine, University of South Dakota), allowing for in-frame replacement of MYC-BioID2 with purified CRY2olig-mCherry. An N-terminal fragment of CRY2oligo-mCherry -SUN2 (AA 1-258) was created by insertion of stop codon using PCR mutagenesis. To create C-terminal fragments of SUN2 (AA 173-738), the CRY2oligo-mCherry-SUN2 construct was mutated to insert a second internal *BSRGI* site using 5’-CAGCAGAGTTCTGTACAGCTCGCGGC-3’ and 5’-GCCGCGAGCTGTACAGAATGCTG-3’primers. Modified plasmids were digested with *BSRGI,* purified and re-ligated. Site-directed mutations were created using the QuikChange XL Site-Directed Mutagenesis Kit (Agilent) according to the manufacturer’s instructions.

MP029-CRY2olig-mCherry plasmid was created by amplifying CRY2oligo-mCherry from Addgene Plasmid #60032 by PCR using 5’-CTGTGCGGCCGCGCCATGAAGATGGACAA-3’ and 5’-CTGTGCCGCTGTACAGCTCGTCCAT-3’ primers, and subsequent cloning into the MP029 vector using the *NotI/KpnI* restriction sites. MP029 is a lentiviral vector with a ubiquitin promoter allowing for moderate to low expression levels, a kind gift from Dr. Murali Palangat (National Cancer Institute, NIH) (Palangat and Larson, 2016). MP029-CRY2-mCherry-SUN2 variants were created by amplifying human full-length SUN2 or SUN2-N-1 (AA 1-260) and SUN2-N-2 (AA 1-226) fragments by PCR from pcDNA3.1^+^/C-(K)DY-SUN2 vector (OHu01874,GenScript # NM_001199579.1) and subsequent cloning into MP029-CRY2-mCherry lentiviral vector using the *NheI/XbaI* restriction sites. Forward PCR primer for full length SUN2, and SUN-N-1 and -N-2 variants was 5-CTGTGCTAGCGCCATGTCCCGAAGAAGCCA-3’ and reverse primers were 5’ CTGTTCTAGAGCCCTAGTGGGCGGGCTCC-3’ for full length SUN2, 5’-CTGTTCTAGAGCTCAGCCCTCATCCGGCCTC-3’ for SUN2-N-1 variant and 5’-CTGTTCTAGACTACAGGCACGTCAGCAAGAG-3’ for SUN2-N-2 variant. The construct consisting of nucleoplasmic and transmembrane domains of SUN2 and the luminal domain of SUN1 was created by amplifying SUN1 C-terminal fragment (AA 245-511) by PCR from pcDNA3.1^+^/C-(K)DY-SUN1 vector (OHu26731,GenScript # NM_001130965.3) using 5’-CTGTTCTAGAATGTTGGCTGGCCGTGG-3’forward and 5’-CTGTCTCGAGGCCCTGACTTGCACGTCCA-3’ reverse primers and subsequent cloning into MP029-CRY2-mCherry-SUN2-N-2 vector using the *XbaI/XhoI* restriction sites. mCherry-DN-KASH plasmid was purchased from Addgene (#125553).

Constructs were transfected into U-2 OS, HEK293T and hTERT-immortalized inducible fibroblasts using Lipofectamine2000 or Lipofectamine LTX with Plus Reagent (Thermo Fisher Scientific) according to manufacturer’s instructions.

### Mice

Animal experimental procedures were carried out according to guidelines in the Guide for the Care and Use of Laboratory Animals from the National Institutes of Health. The mice were fed a chow diet and housed in a virus-free barrier facility with a 12h light and dark cycle. Experimental mice used in this study were HGPS (*G608G^+/+^*) transgenic mice expressing a copy of human progerin (Varga et al., 2006), C57BL/6J mice were used as a control. Mice were sacrificed at 3.5 and 5.5 months of age, and liver, heart, and aorta tissues were collected for analysis. Number of mice used for analysis was as follows: C57BL/6J (liver, n=4; heart, n=4; aorta, n=4, samples were pooled into 1), *G608G^+/+^* (liver, n=12; heart, n=10; aorta, n=15, samples were pooled into 6), *G608G^+/-^* (liver, n=8; heart, n=8; aorta, n=8, samples were pooled into 3). Tissues from a *G609G^+/+^* knock-in mouse strain (liver, n=4-5; heart, n=4-5; aorta, n=4-5, samples were pooled into 2) carrying the HGPS mutation (c.1827C>T;p.Gly609Gly mutation) equivalent to the HGPS c.1824C>T;p.Gly608Gly mutation in the human *LMNA* gene (Osorio et al., 2011) were kindly provided by Carlos Lopez-Otin (University of Oviedo, Spain).

### Immunofluorescence staining

Cells were grown either on glass coverslips (for confocal, SIM and STED microscopy) or on 384-well (Perkin Elmer, 6057300) and 96-well (Matrical, MGB096-1-2-LG-L) microplates for high-throughput imaging, washed once with PBS and fixed for 15 min with 4% paraformaldehyde (PFA). Fixed cells were then washed one time with PBS, permeabilized for 10 min (PBS/0.5% Triton X-100) and washed again. Next, cells were incubated for 1.5 hour with primary antibodies diluted in blocking buffer (PBS, 0.05% Tween20, 5% bovine serum albumin). After two consecutive washes with PBS, cells were incubated for 1 hour with secondary antibodies diluted in blocking buffer and washed once with PBS. SIM samples were mounted in Vectashield antifade mounting media with DAPI (Vector Laboratories), while STED samples were mounted in ProLong glass antifade mountant (Thermo Fisher Scientific). Cells grown on microplates were counterstained with DAPI (Biotium, #40043) or DAPI and HCS CellMask Deep Red Stain (Thermo Fisher Scientific, H32721) for 30 min and washed once with PBS. After washing, cells could be stored for extended periods of time in PBS at 4°C. All steps for IF staining were performed at ambient temperature.

### Immunoprecipitation

hTERT-immortalized doxycycline inducible dermal fibroblasts were lysed in IP buffer (20mM Tris-HCl pH7.5, 150mM NaCl, 2mM EGTA, 2mM MgCl2, 0.5% NP-40, 1mM DTT, 1U/ml Benzonase, 1x Protease inhibitor cocktail set from Milipore) for 1h at 4°C and cleared of insoluble material by centrifugation. 1mg of extract was then incubated with 2μg of α-GRP78/BiP rabbit polyclonal antibody (Abcam, ab21685) or 2μg normal rabbit IgG antibody (Cell Signaling, 2729S) overnight, followed by incubation with Pierce protein A/G magnetic beads (Thermo Fisher Scientific) at 4°C for 4h. Beads were thoroughly washed 3x in IP buffer (w/o Benzonase), and precipitated proteins were eluted using 1x Laemmli sample buffer (Bio-Rad), followed by Western blot analysis. Antibody used for Western blot detection was α-GFP rabbit polyclonal (Abcam, ab290) and α-GRP78/BiP rabbit polyclonal (Abcam, ab21685).

### Western blotting

Total cell lysates were prepared by dissolving ∼750, 000 cells in 150µl 1x Laemmli sample buffer (Bio-Rad) and subsequently denatured for 10 min at 95 °C. Equal amounts of protein extract were loaded onto 4-15% Mini-PROTEAN TGX precast gels (Bio-Rad), separated by SDS-PAGE, and transferred onto PVDF membrane using a Trans-Blot turbo transfer system according to manufacturer instructions (Bio-Rad). Membranes were blocked for 30 min in 5% BSA/TBS-T block buffer (20 mM Tris–HCl pH 7.5, 150 mM NaCl and 0.05% Tween-20) and incubated with primary antibody diluted in 2% BSA/TBS-T overnight at 4°C. Membranes were washed 3x 15 min in TBS-T and incubated with appropriate HRP-conjugated secondary antibodies for 1h at room temperature. Protein detection was performed with the ECL western blotting detection system (Amersham) and imaged using a Bio-Rad ChemiDoc imaging system and ImageLab 6.0.1 software.

### Antibodies

The following antibodies were used for immunofluorescence: α-Hsp110 rabbit polyclonal (StressMarq, SPC-195), α-Hsp90 rat monoclonal (Enzo, ADI-SPA-835), α-Hsc70 rat monoclonal (Abcam, ab19136), α-Hsp40 rabbit polyclonal (StressMarq, SPC-100), α-GRP94 rabbit polyclonal (Abcam, ab3674), α-GRP78/BiP rabbit polyclonal (Abcam, ab21685), α-GRP78 mouse monoclonal (StressMarq, SMC-195), α-PDI P4HB mouse monoclonal (Abcam, ab2792), α-Calnexin rabbit polyclonal (Cell Signaling, 2433S), α-SUN1 rabbit polyclonal (Sigma, HPA008346), α-SUN2 rabbit polyclonal (Sigma, HPA001209) and α-Nucleoporin rat monoclonal (Abcam, ab188413). For western blotting the following antibodies were used: α-Hsp110 rabbit polyclonal (StressMarq, SPC-195), α-Hsp90 rat monoclonal (Enzo, ADI-SPA-835), α-Hsc70 rat monoclonal (Abcam, ab19136), α-Hsp40 rabbit polyclonal (StressMarq, SPC-100), α-IRE1 (phospho S724) rabbit monoclonal (Abcam, ab124945), α-IRE1 rabbit polyclonal (Abcam, ab37073), α-Phospho-PERK (Thr982) rabbit monoclonal (Abcam, ab192591), α-SUN1 rabbit polyclonal (Sigma, HPA008346), α-SUN2 rabbit polyclonal (Sigma, HPA001209), α-GFP rabbit polyclonal (Abcam, ab290) and α-GAPDH mouse monoclonal (Abcam, ab8245). Secondary antibodies used for immunofluorescence detection in confocal and SIM microscopy were Alexa Fluor Rabbit-anti-Mouse 568 (Invitrogen, A11004) and Donkey-anti-Mouse 647 (Invitrogen, A31571), Alexa Fluor Donkey-anti-Rabbit 568 and 647 (Invitrogen, A10042 and A31573), Alexa Fluor Donkey-anti-Rat 568 (Invitrogen, A11077) and Chicken-anti-Rat 647 (Invitrogen, A21472). Secondary antibodies used for STED detection were Abberior Goat-anti-Rabbit STAR RED (Sigma, 41699) and AffiniPure Fab Fragment Goat Anti-Mouse Alexa Fluor 594 (Jackson Immunoresearch, 115-587-003). Secondary antibodies used for western blot were Mouse-anti-rabbit IgG-HRP (Santa Cruz, sc-2357), Goat-anti-Mouse IgG-HRP (Santa Cruz, sc-2005) and Goat-anti-Rat IgG-HRP (Santa Cruz, sc-2032). All antibodies were used at dilutions recommended by the manufacturer.

### RNA isolation and quantitative analysis of gene expression

Total RNA was extracted from cells using the NucleoSpin RNA Kit (Takara Bio) according to manufacturer instructions. Total RNA from flash frozen liver, heart and aorta tissues was isolated from 25-50 mg ground tissue. Briefly, tissue was homogenized in 600 μl lysis buffer using a Bullet Blender, 0.9-2.0 mm stainless steel beads for 3 mins, setting 12 at 4°C, total RNA was isolated using RNeasy Mini Kit (Qiagen) and the integrity of the RNA was assessed by Bioanalyzer using RNA 6000 Nano Chip (Agilent, Inc.). mRNA levels of UPR genes in control and progerin-expressing cells and mice were measured by reverse transcribing 1 μg of RNA using iScript cDNA synthesis kit containing a blend of oligo(dT), random hexamer primers and iScript™ Reverse Transcriptase according to manufacturer recommendation (Biorad). Equal volumes of cDNA were used as template in a real-time quantitative PCR reaction using iQ SYBR Green Supermix (BioRad) on a CFX384 Touch Real-Time PCR System (BioRad). Reaction conditions were: 3 min at 95°C, 1 cycle; 20 s at 95°C, 30 s at 58°C, 40 cycles. Melting curves of the amplified product were generated to verify that a single amplicon was generated. All the values were normalized to the internal controls glyceraldehyde 3-phosphate dehydrogenase (GAPDH) and TATA-box-binding protein 1 (TBP1) genes. Primer combinations are indicated in Table S2. Expression values are shown as means ± s.d. of biological triplicates and statistical significance between biological replicates was determined by Student t-test in Graphpad Prism and Excel.

### siRNA Transfection

For siRNA treatment hTERT immortalized GFP-progerin doxycycline inducible dermal fibroblasts were plated on a 10 cm^2^ culture well (6-well) at 70-80% confluence in high glucose MEM containing 15% FCS and 0.2mM L-glutamine 16 h prior to transfection. Cells were transfected on two consecutive days with each siRNA (working concentration of 10nM) and 5μl Lipofectamine RNAiMAX (Thermo Fisher Scientific) according to manufacturer instructions. After 96h, cells were collected and plated onto 384-well plates for high-throughput imaging analysis. Silencing efficiency of each protein was monitored by western blot analysis and immunofluorescence. All experiments were performed in triplicates. siRNAs used were Silencer Select negative control (Ambion, #4390843), Silencer Select UNC84A siRNA (Ambion, #4392420 and custom UNC84B siRNA 5’-CGUACCAGGUGGUGGAGCUGCGGAU-3’ (Dharmacon).

### High-Throughput Image Acquisition and Analysis

Cells were imaged in four channels (405, 488, 561, and 640 nm excitation lasers) in an automated fashion using a dual spinning disk high-throughput confocal microscope (Yokogawa CV7000) with a 405/488/561/640 nm excitation dichroic mirror, a 40 air objective lens (NA 0.95) or a 60X water immersion lens (NA = 1.2),, a 568 nm excitation dichroic mirror, and two 16-bit sCMOS cameras with binning set to 2 (Pixel size: 0.325 nm for 40X and 0.216 nm for 60X nm). Emission bandpass filters were used for each channel: 445/45, 525/50, 600/37, and 690/29 nm. Nine randomly selected fields of view were imaged per well in a single optimal focal plane. Images were corrected on the fly using a geometric correction for camera background, illumination field (vignetting), camera alignment, and chromatic aberrations. Corrected images were saved and stored as 16-bit TIFF files. Captured images were then analyzed using Columbus 2.8.1 or 2.9.1 (Perkin Elmer). Briefly, nuclei regions of interest (ROI) were segmented using the DAPI (405 nm) channel and cytoplasmic based ROIs were segmented using the CellMask HCS channel (640 nm). Cytoplasm and nucleus ROIs adjacent to the image edges were excluded from subsequent image analysis steps. The mean fluorescence intensity for the nucleus and cytoplasmic ROI was measured in the 561 nm channel. Analysis of the recruitment of GRP78/BiP to the nuclear periphery was done by creating a ring ROI covering 5-35% of the distance from the DAPI-based nuclear ROI (where 0% is adjacent to DAPI and 100% represents the cell border) and the mean fluorescence intensity per well for ring ROI was measured in 561nm channel. Columbus results were exported as comma-separated text files and analyzed using R. 300-1500 cells were analyzed per condition. Data are plotted as means of 3 biological replicates +/-standard deviation (SD).

### Light Microscopy

Laser scanning confocal microscopy was performed using a Carl Zeiss LSM880 inverted microscope controlled by ZEN software with definite focus. Cells were imaged using Plan Apochromat 63×/1.46 NA oil objective and images were acquired in three channels (405, 488 and 561nm excitation lasers). Raw images were reconstructed using the appropriate tool in the Zeiss Zen software and ImageJ. *In vivo* FRAP experiments were performed using a Carl Zeiss LSM780 inverted microscope controlled by ZEN software and with definite focus. Live cells were imaged using 63×/1.46 NA oil objective and a CO_2_/heating control stage insert. Induced MP029-CRY2olig-mCHerry-SUN2-N clusters were bleached using 488 and 561 nm light with maximum laser power and recovery was monitored over a period of 2–5 min.

### Super Resolution Microscopy and Image Quantification

Structured Illumination Microscopy was performed on fixed cells using a Zeiss Elyra PS.1 on an AxioObserver Z1 inverted microscope controlled by Zen software (Carl Zeiss). Images were acquired in four channels (405, 488, 561, and 640 nm excitation lasers) using Plan Apochromat 63x/1.4 NA oil immersion objective (Carl Zeiss) and Pco.edge sCMOS camera in a 30°C environmental chamber. Raw images were reconstructed using the appropriate tool in the Zeiss Zen software (Black version with Structured Illumination module). Stimulated Emission Depleted (STED) microscopy was performed on fixed cells using a commercial Leica SP8 STED 3X system (Leica Microsystems), equipped with a white light laser (range, 470–670 nm) and 592-nm, 660-nm and pulsed 775-nm STED depletion lasers. Special STED white objective 100×/1.4-NA oil-immersion objective lens (HCX PL APO STED white, Leica Microsystems) was used for imaging and sequential confocal and STED images were taken in three channels via three sequential excitations. GFP labelled probes were excited with the 488 nm white light laser and depleted with a 592 nm STED laser at 25% power, Alexa Fluor 594 probes were excited with the 570 nm wavelength white light laser and depleted with a 775 nm STED laser at 20% power and Abberior STAR RED probes were excited with the 647 nm white light laser and depleted with a 775 nm STED laser at 20% power. All images were acquired in 2D STED mode with a Z-stack size of 0.12μm, a scan speed of 600 lines per second, a pixel size of 50–80 nm (512× 512 pixels) and 8 lines averages. Deconvolution of STED data was carried out using the STED module in Huygens Professional software version 14.10.1 (Scientific Volume Imaging) using the classical maximum likelihood estimation algorithm and a STED saturation factor of 9.

Data was reconstructed with Fiji software and images were montaged with Adobe Photoshop CC 2019 software. Fluorescence intensity values in line scans were generated using the corresponding tools in Fiji or Zeiss Zen software. Local enrichment of SUN2, GRP78 and PDI was measured by drawing ROI at the nuclear periphery using the corresponding tools in Fiji or Zeiss Zen software.

### Live Cell Imaging and Protein Clustering Induction

96-well plates (Matrical, MGB096-1-2-LG-L) were coated for 1h with 0.1% gelatin solution (ATCC, PCS-999-027) and washed twice with PBS (pH 7.4, Thermo Fisher Scientific). HEK293FT cells were plated on the gelatin coated dish and grown overnight in normal growth medium to reach ∼ 70% confluency. Constructs were transfected on the next day using Lipofectamine2000 (Thermo Fisher Scientific) according to manufacturer instructions and live cell imaging was performed 24h after the transfection. Just prior to imaging, the medium was replaced with 150μl of imaging medium (Phenol Red free DMEM medium containing 15% fetal bovine serum). Live-cell high-throughput imaging experiments were performed using a Yokogawa CV8000 high-throughput dual spinning disk confocal microscope with 60x water immersion lens (NA = 1.2). Cells were incubated in an environmental chamber at 37°C, 5% CO_2_ and 80% humidity. For protein clustering induction, cells were imaged by use of two laser wavelengths (488 nm for CRY2 activation /560 nm for mCherry imaging). To execute CRY2activation, the repetitive ON/OFF cycle was applied every 10 min for 90 min (488nm activation duration was fixed to 200ms with 60% laser power in all measurements). Image z-stacks of 5 planes at a 1μm interval were acquired using a heated Olympus PlanApoChromat 60X water lens objective (NA 1.2), and 2 sCMOS cameras (2550 X 2160 pixels) using camera binning of 2X2. A total of 16 fields of view were acquired. Images were corrected on the fly using a geometric correction for camera alignment and optical aberrations and saved and stored as 16-bit TIFF files. Data was reconstructed with Fiji software and images were montaged with Adobe Photoshop CC 2019 software.

### Cell Sorting and Protein Clustering Induction on a 24-well plate

For fluorescence-activated cell sorting (FACS), cells were transfected with MP029-CRY2olig-mCherry, MP029-CRY2olig-mCh-SUN2-N-1 or MP029-CRY2olig-mCh-SUN2-N-2 using Lipofectamine LTX with Plus Reagent (Thermo Fisher Scientific) according to manufacturer instructions. After 24h cells were trypsinized, collected in PBS and examined for mCherry expression using a BD FACSAria Cell Sorting System with BD FACSDiva Software (BD Biosciences). Non-fluorescent cells were negatively selected and the remaining gated cells were sorted based on the presence of mCherry fluorescent protein. Sorted cells were plated in a 24-well plate at a density of 5-6×10^5^ cells/well for RNA isolation or on a 8-well microscopy chamber at a density of 1×10^5^ cells/well for immunofluorescence and left to recover overnight. The next day CRY2 activation was executed using a UV LED light source (M405L4, ThorLabs) operated by LED Driver (DC2200, ThorLabs). Repetitive ON/OFF cycle was applied every 10 sec for 60 min (405nm activation duration was fixed to 200ms with 5% power in all measurements). Cells were then either fixed with 4% PFA for immunofluorescence analysis or collected for RNA isolation and subsequent RT-qPCR analysis as described above.

### Statistical Analysis

High-throughput imaging experiments in inducible cell lines were performed on three technical replicates (wells) and averaged to obtain the measurement for that experiment. The experiments were repeated independently three times (*n* = 3), and the values for each experiment were averaged. Data calculations were performed using R software and paired two-tailed Student’s t-test for the fold change (FC) indicated in the figures was used for statistical analyses. High-throughput imaging experiments in immortalized and primary patient-derived fibroblasts were performed on three technical replicates (wells) and the experiments were repeated three times (*n* = 3) for subsequent passage numbers. Single cell data analysis was performed using R software and the data are shown as the average mean values of 3 different experiments and/or the percentage of cells showing increase in the phenotype relative to the wild-type control. The threshold for quantification of increases was set as mean of the wild-type +/-1 SD. Statistical analysis for the average mean values of single cell data was performed by an ANOVA one-way test, followed by a post-hoc Dunnett’s test using the cell line as the predictor variable and fixing WT as the negative control. Statistical analysis to determine any difference between cell lines for the percentage of cells showing increase or decrease was performed by Chi-square test. For FRAP experiments, all FRAP measurements were pooled together and averaged (n=20). Fitted FRAP curve was obtained by fitting the recovery portion to the equation R = C + P(1 – exp(-kt). All other calculations were performed using Graphpad Prism software and Excel and paired two-tailed Student’s t-test was used for statistical analyses. Error bars represent standard deviation (s.d.) or or standard error of the mean (s.e.m.) as indicated. Statistical significance was classified as follows: *p<0,05, **p<0,005, ***p<0,0005.

## AUTHOR CONTRIBUTIONS

S.V. and T.M. designed the study; L.S. designed CRY2oligo-mCherry clustering experiments, generated CRY2oligo-mCherry-SUN2 full length and deletion constructs and optimized clustering conditions; G.P. wrote the R code and performed single cell analysis high-throughput image in HGPS patient-derived cells. S.V. performed all other experiments and data analysis, S.V and T.M. wrote the manuscript. All authors read and approved the manuscript.

## ACKNOWLEDGMENTS

We thank the Misteli lab members for sharing feedback, data, and reagents. We are grateful to Susan Shackleton (University of Leicester, UK), Kyle Roux (Sanford School of Medicine, University of South Dakota), Chandra Tucker (University of Colorado School of Medicine) and Murali Palangat (National Cancer Institute, NIH) for providing plasmids, Francis Collins (National Human Genome Research Institute, NIH) for providing HGPS G608G mice, Carlos Lopez-Otin (University of Oviedo, Spain) for providing tissue of HGPS G609G mice, Tatiana Karpova and David Ball for help with Structured Illumination Microscopy, Laser Scanning Confocal Microscopy and FRAP experiments as part of the NIH/NCI/CCR LRBGE Optical Imaging Core, NIH/NCI/CCR High Throughput Imaging Facility (HiTIF) for help with high throughput imaging and automated liquid handling, and Daniela Malide and Christian Combs at the NIH/NHLBI Light Microscopy Core for help with STED imaging. The T.M. lab is supported by the Intramural Research Program of the NIH, NCI, Center for Cancer Research and S.V. was supported by FWF Austrian Science Fund (Vienna), GRANT NUMBER: J 3849.

---

**Figure S1.**
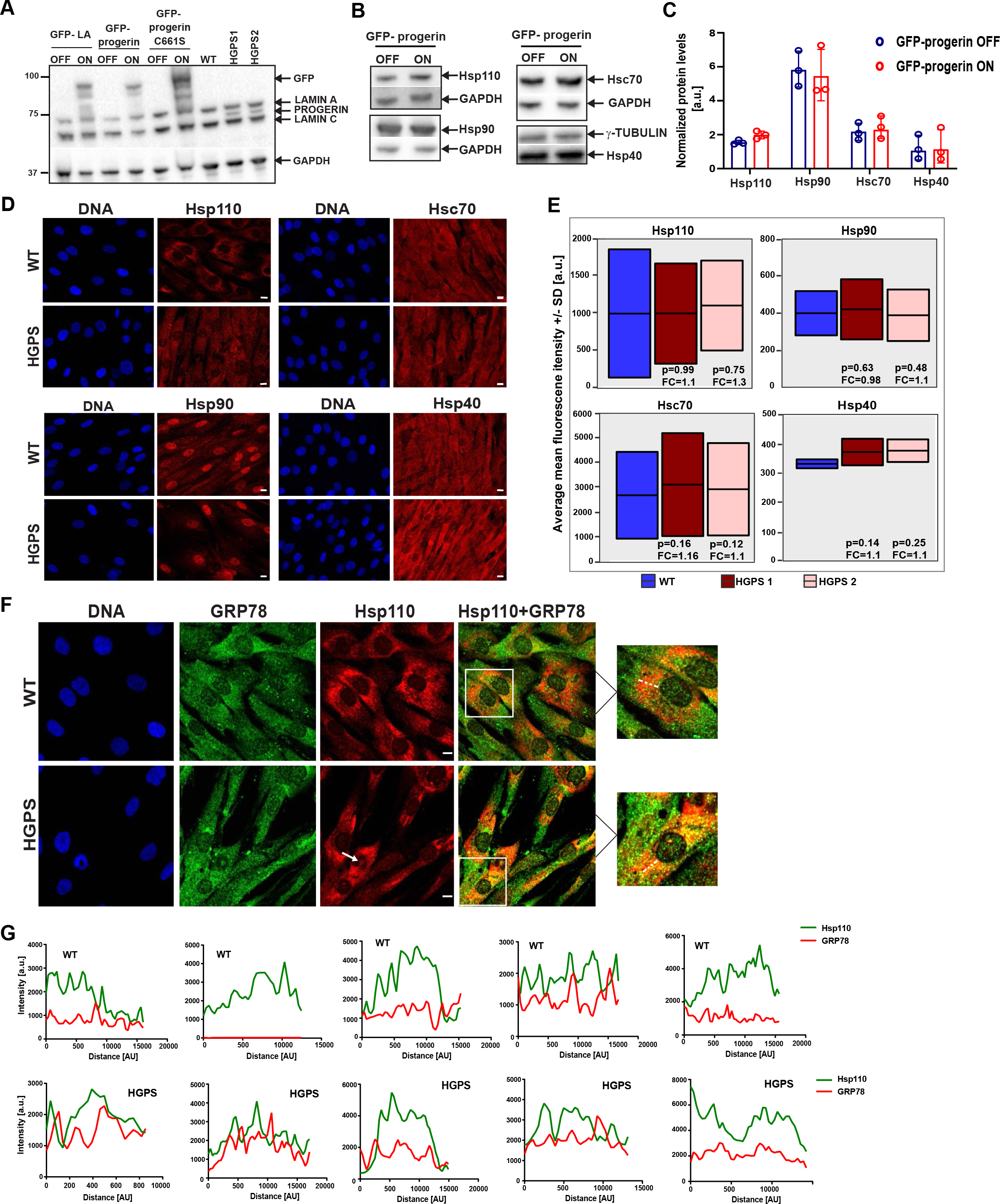
Levels of cytosolic chaperones in progerin-expressing cells. **(A)** Western blot analysis of total protein extract from inducible and immortalized patient-derived fibroblasts using anti-lamin A/C antibody to detect endogenous lamin A/C, progerin and GFP-tagged variants. GAPDH served as a loading control. **(B)** Immunoblot analysis of cytosolic chaperones in total cell lysates of GFP-progerin inducible fibroblasts. GAPDH served as a loading control. **(C)** The protein levels in Fig. S1A normalized to housekeeping GAPDH gene and shown as mean protein levels of 3 different experiments. **(D)** IF staining of several cytosolic chaperones in PFA fixed patient derived HGPS fibroblasts and their respective control. Scale bar: 10μm. **(E)** High throughput IF quantification of cytosolic chaperone expression levels in patient-derived fibroblasts represented as the average mean fluorescence values +/-SD from 3 experiments (n=300-500). Statistical differences were analyzed by *t*-test for the fold change indicated in the figure. **(F)** Representative IF staining for Hsp110 and GRP78 in PFA fixed control fibroblasts (WT) and HGPS patient-derived fibroblasts. Arrows indicate Hsp110 accumulation in progerin expressing cells. Scale bar: 10μm. **(G)** Line graphs indicate IF signal intensities along a dotted line based on an overlap in 488nm and 594nm channels in 5 randomly selected cells.

**Figure S2.**
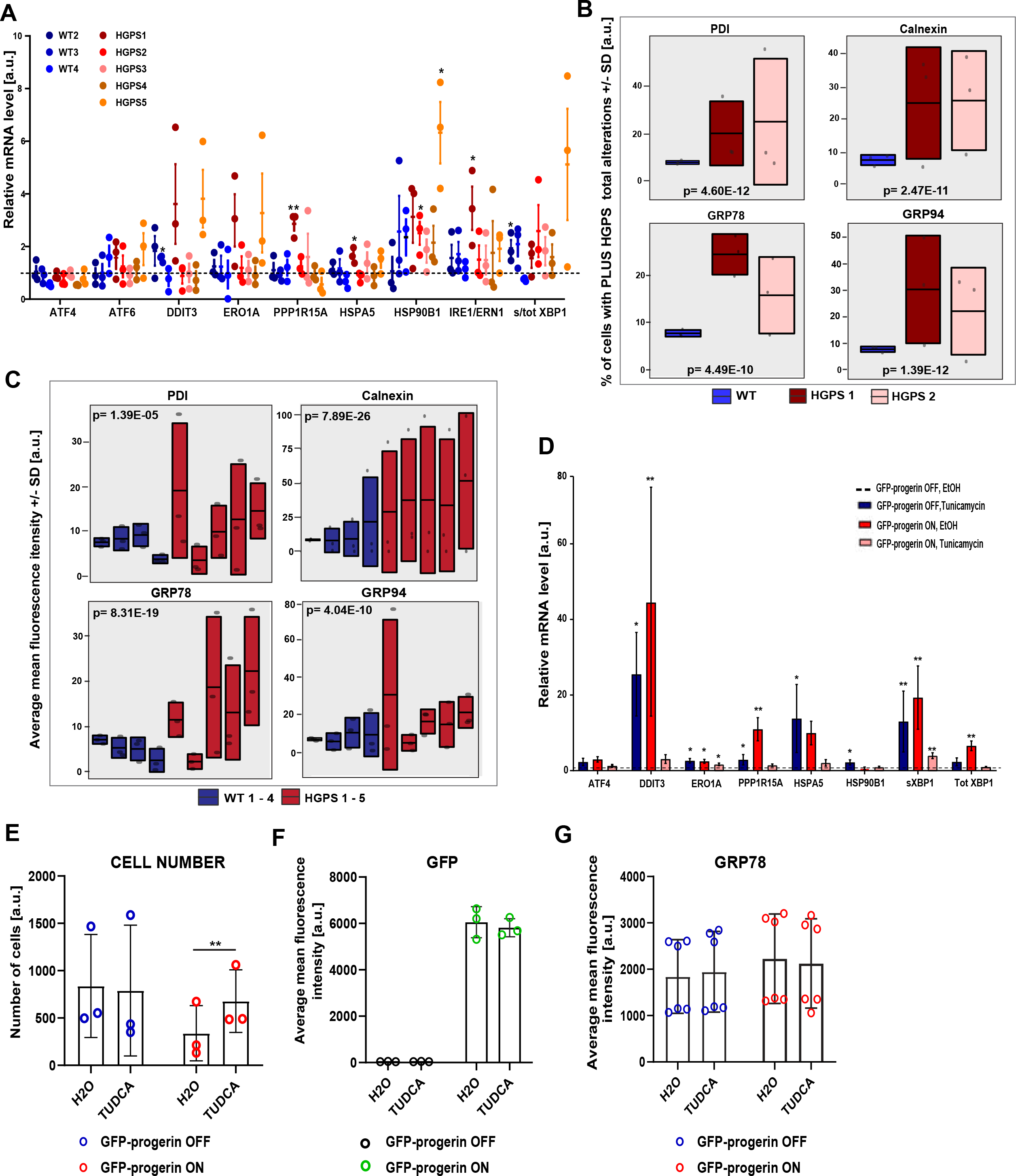
Chronic activation of ER stress in progerin-expressing cells. **(A)** Single cell high throughput quantification of ER chaperone expression levels in immortalized human HGPS fibroblasts (n=300) represented as the % of cells showing an increase in the mean intensity level +/-1 SD relative to the control (WT) in 3 biological replicates (See Experimental Procedures). Statistical differences were analyzed by Chi-square test**. (B)** Single cell high throughput quantification of ER chaperone expression levels in five patient-derived primary human HGPS fibroblasts *HGPS 1-5) and 4 healthy controls (WT 1-4) represented as the % of cells showing an increase in the mean intensity level +/-1 SD relative to the control (WT) in 3 biological replicates (n=50-300). Statistical differences were analyzed by Chi-square test**. (C)** Quantification of mRNA levels of UPR genes in five patient-derived primary human HGPS fibroblasts and 4 healthy controls relative to WT 1. *GAPDH* and *TBP1* were used for normalization. Values represent the mean of 3 biological replicates ± SD. Statistical differences were analyzed by *t*-test for HGPS 1-5/WT 1 and WT 2-4/WT 1 \**P*<0.05. Dashed lines represent control value (WT 1) normalized to 1. **(D)** Quantification of mRNA levels of several UPR genes in GFP-progerin inducible fibroblasts treated with Tunicamycin or EtOH (vehicle control) by qRT-PCR. Values for GFP-progerin OFF/Tunicamycin and GFP-progerin ON/EtOH were calculated relative to GFP-progerin OFF/EtOH control, whereas GFP-progerin ON/Tunicamycin is shown relative to GFP-progerin ON/EtOH. *GAPDH* and *TBP1* were used for normalization. Values represent means of 3 biological replicates ± SEM. Dashed line represent control normalized to 1. Statistical differences were analyzed by *t*-test. \**P*<0.05, \*\**P*<0.01. **(E)** Number of cells in GFP-progerin inducible fibroblasts treated with 25uM TUDCA or H2O (vehicle control) for 48h. Values represent means of 3 biological replicates ± SD (n=350-1550). Statistical differences were analyzed by *t*-test. \**P*<0.05, \*\**P*<0.01. **(F)** Average mean GFP-progerin levels in inducible fibroblasts treated with 25uM TUDCA or H2O (vehicle control) for 48h. Values represent means of 3 biological replicates ± SD (n=350-1550). Statistical differences were analyzed by *t*-test. **(G)** Average mean GRP78 levels in inducible fibroblasts treated with 25uM TUDCA or H2O (vehicle control) for 48h. Values represent means of 3 biological replicates ± SD (n=350-1550). Statistical differences were analyzed by *t*-test.

**Figure S3.**
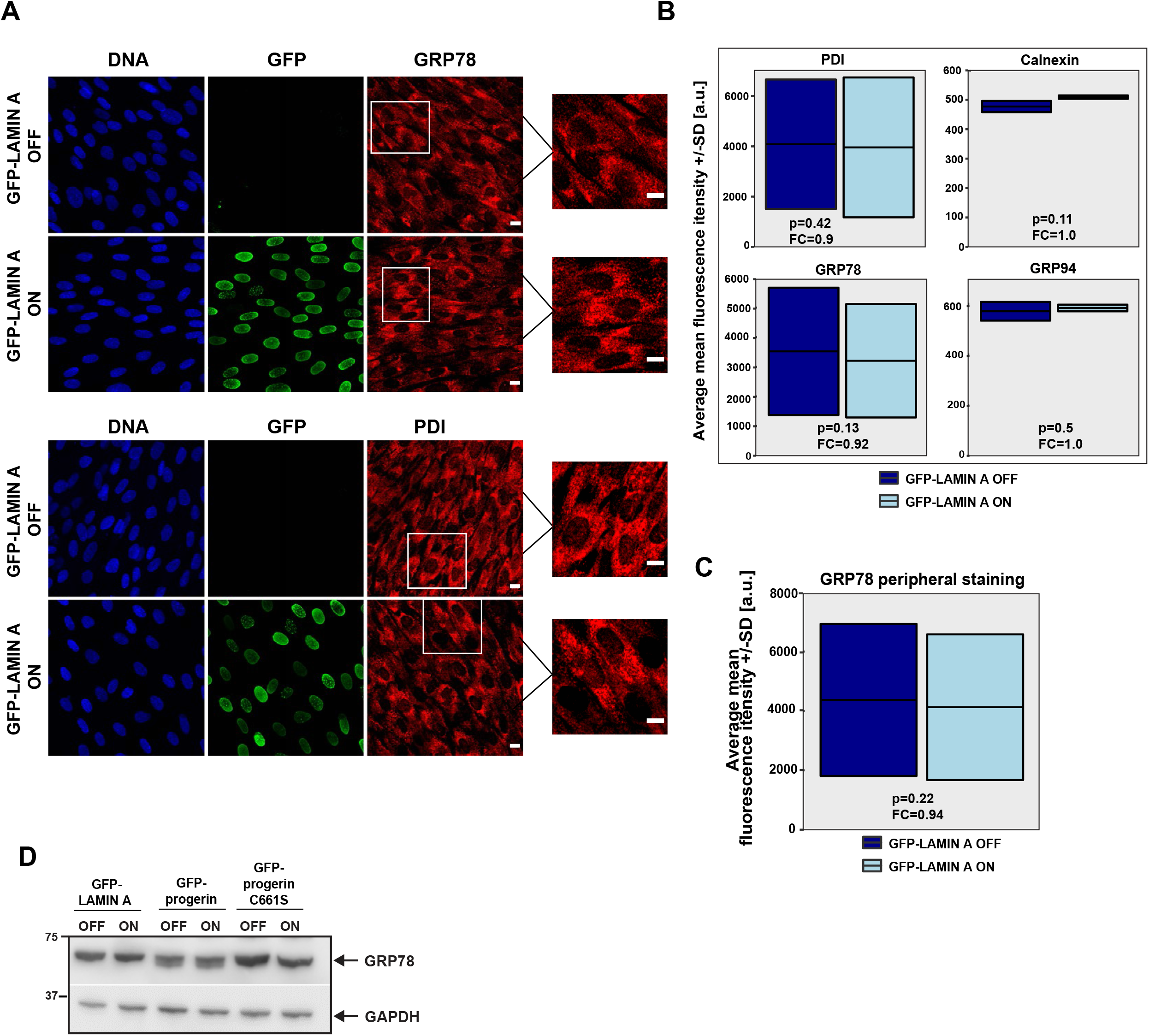
Expression of wild type GFP-lamin A does not induce ER stress. **(A)** Representative confocal images of GRP78 and PDI staining in GFP-lamin A inducible fibroblasts fixed with PFA. Scale bar: 10μm. **(B)** High throughput IF quantification of ER chaperone expression levels in GFP-lamin A inducible fibroblasts represented as the average mean fluorescence intensity +/-SD from 3 different experiments (n=1000) (See Experimental Procedures). Statistical differences were analyzed by *t*-test for the fold change indicated in the figure. **(C)** High throughput IF quantification of GRP78 recruitment to the nuclear periphery in GFP-lamin A inducible fibroblasts (n=1000) represented as the average mean fluorescence intensity +/-SD from 3 experiments. Statistical differences were analyzed by *t*-test for the fold change indicated in the figure. **(D)** Immunoblot analysis of GRP78 levels in total cell lysates of GFP-lamin A, GFP-progerin and GFP-progerinC661S fibroblasts used for immunoprecipitation. GAPDH served as a loading control.

**Figure S4.**
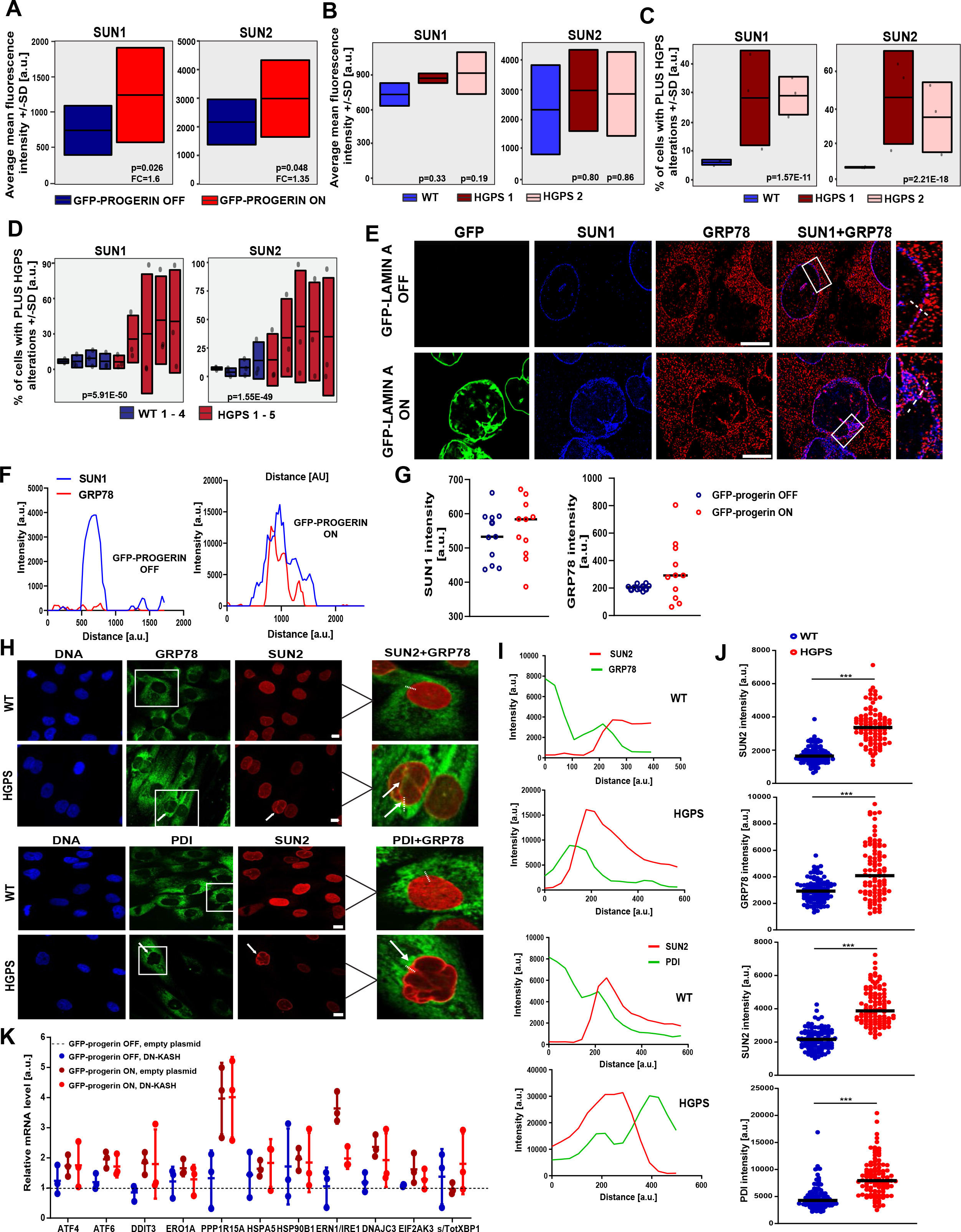
SUN1 and SUN2 levels are increased in progerin-expressing cells. **(A)** High throughput IF quantification of SUN1 and SUN2 expression levels in GFP-progerin inducible fibroblasts (n=1300-1500) represented as the average mean fluorescence intensity +/-SD from 3 experiments (See Experimental Procedures). Statistical differences were analyzed by *t*-test for the fold change indicated in the figure. **(B, C)** Single cell high throughput IF quantification of SUN1 and SUN2 expression levels from 3 experiments in immortalized human HGPS fibroblasts (n=200-400) represented as the distribution of average mean fluorescence intensity +/-SD **(B)** and the % of cells showing an increase in the mean intensity level +/-1 SD relative to the control (WT) **(C)** (See Experimental Procedures). Statistical differences were analyzed by Dunnett’s test**. (D)** Representative SIM images of GRP78 and SUN1 staining fixed with PFA. Scale bar: 10μm. **(E)** Line graphs indicating IF signal intensity across the dotted lines in panel C based on an overlap in 594nm and 647nm channels. **(F)** Scatter plots showing fluorescence intensities of the peripheral SUN1 and GRP78 (n=11) in SIM images of GFP-progerin inducible fibroblasts. Line represents the median. Statistical differences were analyzed by *t*-test. **(G)** Representative confocal images of GRP78, PDI and SUN2 staining in patient-derived fibroblasts fixed with PFA. Scale bar: 10μm. Arrows indicate GRP78 recruitment to the nuclear periphery. **(H)** Line graphs indicating IF signal intensity across the dotted lines in panel G. **(I)** Scatter plots showing Fluorescence intensities of the peripheral SUN2, GRP78 and PDI (n=100) in immortalized patient-derived fibroblasts shown as scatterplots. Line represents the median. Statistical differences were analyzed by *t*-test. ***P<0.005. **(J)** Quantification of mRNA levels of UPR genes in GFP-progerin cells transfected with DN-KASH or an empty plasmid relative to the uninduced empty plasmid control. *GAPDH* and *TBP1* were used for normalization. Values represent the mean of 3 biological replicates ± SD. Dashed lines represent control value normalized to 1.

**Figure S5.**
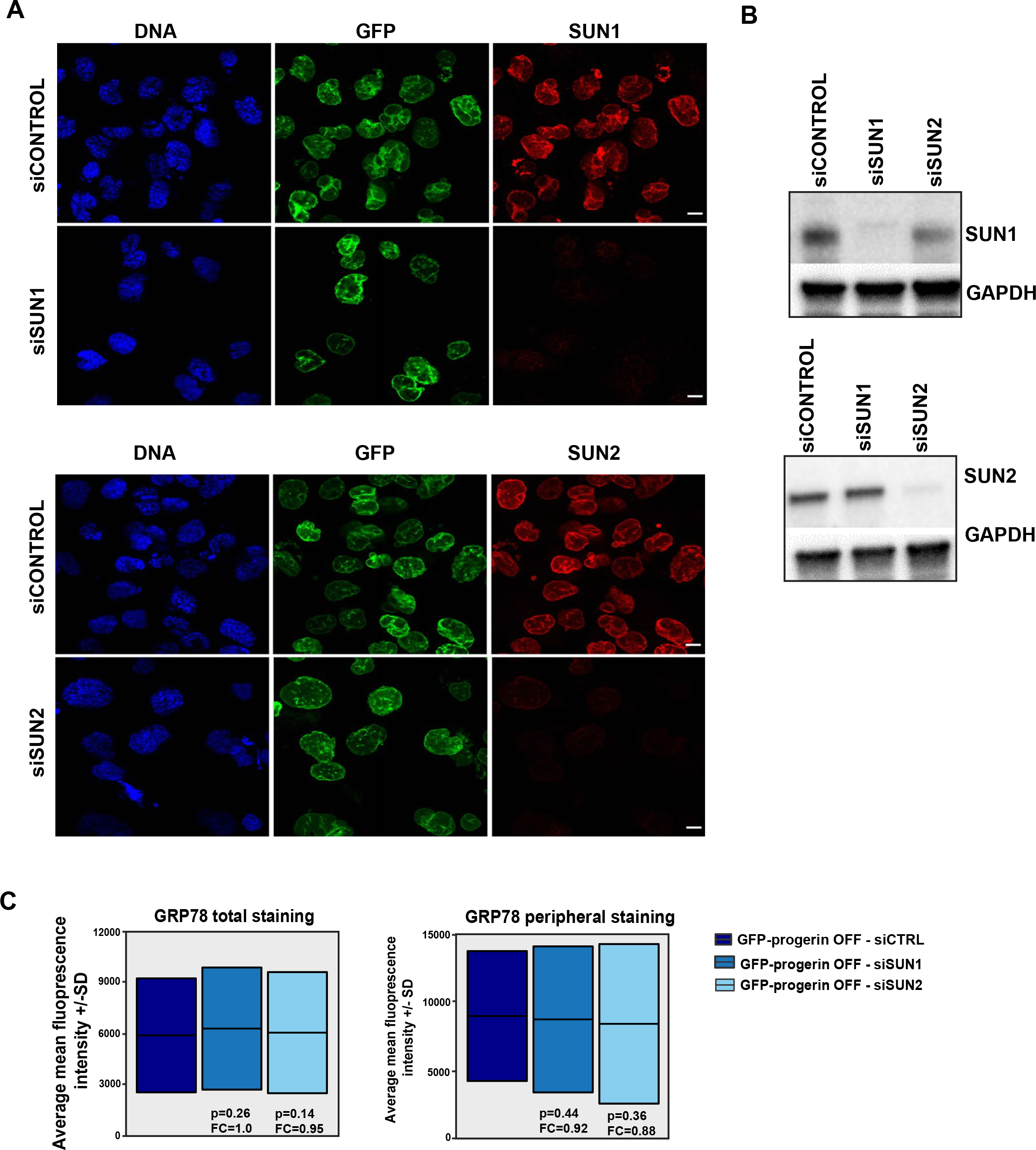
SUN1 and SUN2 are efficiently downregulated in GFP-progerin cells. **(A)** Representative confocal images of SUN1 and SUN2 staining in siSUN1 and siSUN2 treated GFP-progerin inducible fibroblasts and their respective controls. Scale bar: 10μm. **(B)** Immunoblot analysis of SUN1 and SUN2 expression in total cell lysates of siSUN1 and siSUN2 treated GFP-progerin fibroblasts and their respective controls. GAPDH served as a loading control. **(C)** High throughput IF quantification of the total GRP78 levels and the GRP78 recruitment to the nuclear periphery in siSUN1 and siSUN2 treated GFP-progerin OFF fibroblasts (n = 200-400) represented as the average mean fluorescence values +/-SD from 3 experiments. Statistical differences were analyzed by *t*-test for the fold change (FC) indicated in the figure.

**Figure S6.**
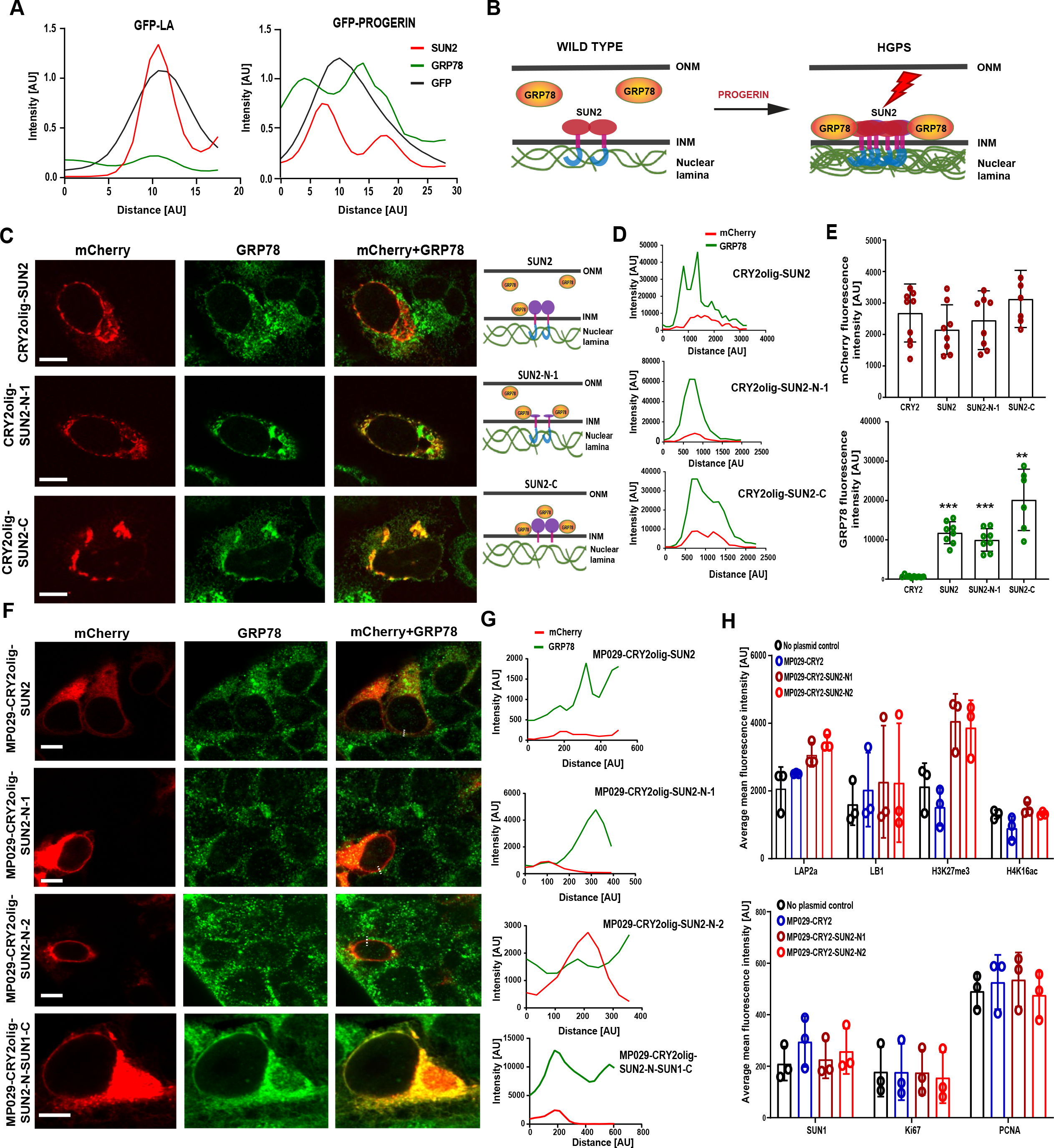
Expression and clustering of SUN2 variants in wild type cells. **(A)** Line graphs indicating IF signal intensity across the dotted lines in Fig. 5A. Data were normalized to the highest GFP signal (n=20). **(B)** Mechanistic model of SUN2-induced ER stress response. Progerin aggregates at the nuclear periphery cause SUN2 clustering in the INM via its interaction with the nucleoplasmic SUN2 N-terminus. SUN2 clusters are sensed in the ER lumen by GRP78/BiP through the SUN2 C-terminal domain, leading to chronic ER stress and UPR activation. **(C)** Representative confocal images of CRY2olig-mCherry SUN2 variants after CMV-promoter driven high level expression and GRP78 staining in U2OS cells fixed with PFA. Scale bar: 10μm. Arrows indicate SUN2 and GRP78 clusters. Schematic drawings on the right depict GRP78 recruitment by various SUN2 variants in the INM. **(D)** Line graphs indicating IF signal intensity across the clusters (dashed lines) based on an overlap in 488nm and 594nm channels in panel D. (E) Mean fluorescence intensities of the mCherry and GRP78 in U2OS cells transfected with various CRY2olig-mCherry constructs (n=6-9). Statistical differences were analyzed by *t*-test relative to CRY2olig only control. \*\**P*<0.005, ***P<0.0005. **(F)** Representative confocal images of GRP78 staining in HEK293FT cells expressing either full length MP029-CRY2-mCherry-SUN2 or N-terminus variants depicted in Fig. 5C. Scale bar: 10μm. **(G)** Line graphs indicating IF signal intensity across the clusters (dashed lines) in panel C. **(H)** Average mean fluorescence intensity levels of several HGPS cellular and proliferation markers in HEK293FT cells transfected with various MP029-CRY2-mCherry constructs.

**Figure S7.**
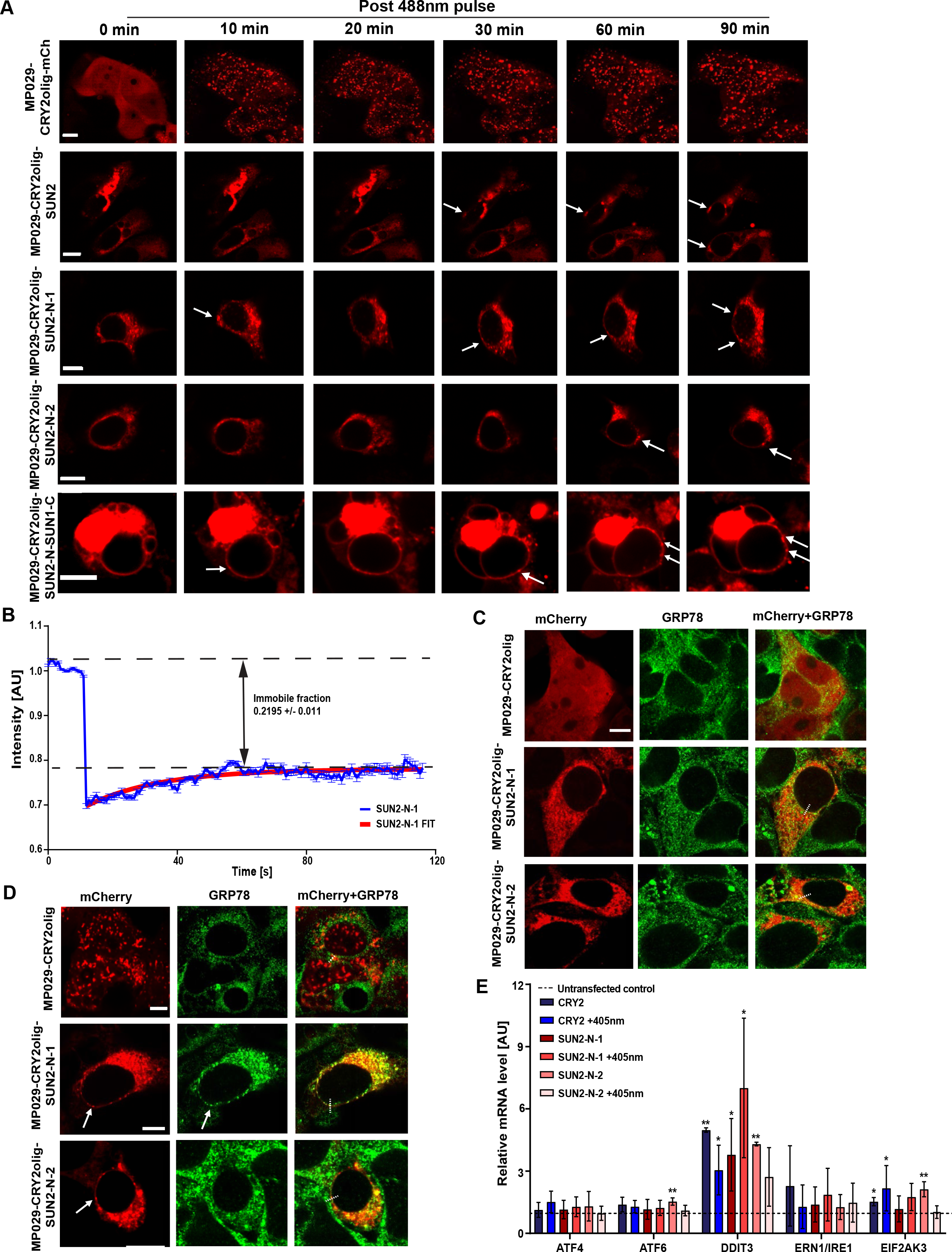
Expression and clustering of SUN2 variants in HE293FT cells after cell sorting. **(A)** Live cell imaging of HEK293FTcells expressing MP029-CRY2olig-mCherry variants pre and post blue light (200ms pulse, 488nm, 60% laser power every 10 min for 90 min). Scale bar: 10μm. Arrows indicate clusters. **(B)** Representative confocal images of GRP78 staining in HEK293FT cells expressing MP029-CRY2olig-mCherry or MP029-CRY2olig-mCherry SUN2 N-terminus variants after the cell sorting. Scale bar: 10μm. **(C)** Representative confocal images of GRP78 staining in HEK293FT cells expressing MP029-CRY2olig-mCherry or MP029-CRY2olig-mCherry SUN2 N-terminus variants after cell sorting and blue light exposure (200ms pulse, 405nm, 5% power every 10 sec for 60 min). Scale bar: 10μm. Arrows indicate clusters. **(D)** Fluorescence recovery curve of FRAP analysis of formed clusters in HEK293FT cells expressing MP029-CRY2olig-mCherry SUN2 N-terminus variant after blue light exposure. Data represent mean ± s.e.m. (n=20). (**E**) Quantification of mRNA levels of several UPR genes in sorted HEK293FT cells expressing MP029-CRY2olig-mCherry or MP029-CRY2olig-mCherry SUN2 N-terminus variants before or after blue light exposure relative to untransfected control. *GAPDH* and *TBP1* were used for normalization. Values represent the mean of 3 biological replicates ± SD. Statistical differences were analyzed by *t*-test. \**P*<0.05, \*\**P*<0.01. Dashed lines represent control value normalized to 1.

**Figure S8.**
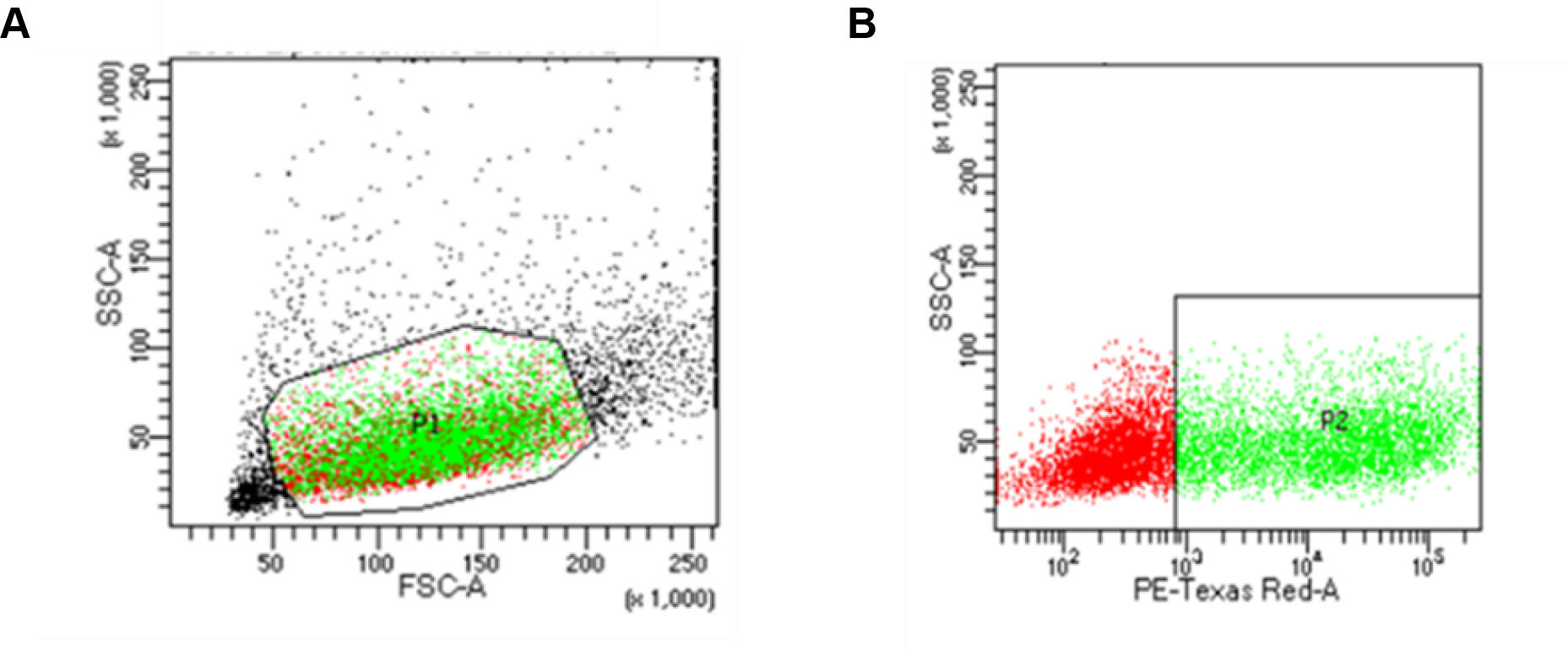
Cell sorting strategy. **(A)** Cell size (forward scattering area, FSC-A) and granularity (side scatter area, SSC-A) were used to identify the main cell fraction (P1). (B) Non-fluorescent cells were negatively selected and the remaining cells were gated based on the presence of mCherry fluorescent protein (P2).

**Table S1.** Primary patient-derived cell lines and healthy controls.

| **Cell line** | **Affected** | **Age at sampling** | **Source** |
| --- | --- | --- | --- |
| GM00038 (WT 1) | No | 9 years | Coriell |
| AG08470 (WT 2) | No | 10 years | Coriell |
| HGMDFN090 (WT 3) | No | 37 years | PRF |
| HGFDFN168 (WT 4) | No | 40 years | PRF |
| AG08466 (HGPS 1) | Yes | 8 years | Coriell |
| HGADFN127 (HGPS 2) | Yes | 3 years | PRF |
| GM01972D (HGPS 3) | Yes | 14 years | Coriell |
| HGADFN167 (HGPS 4) | Yes | 8 years | PRF |
| HGADFN178 (HGPS 5) | Yes | 6 years | PRF |

**Table S2.**
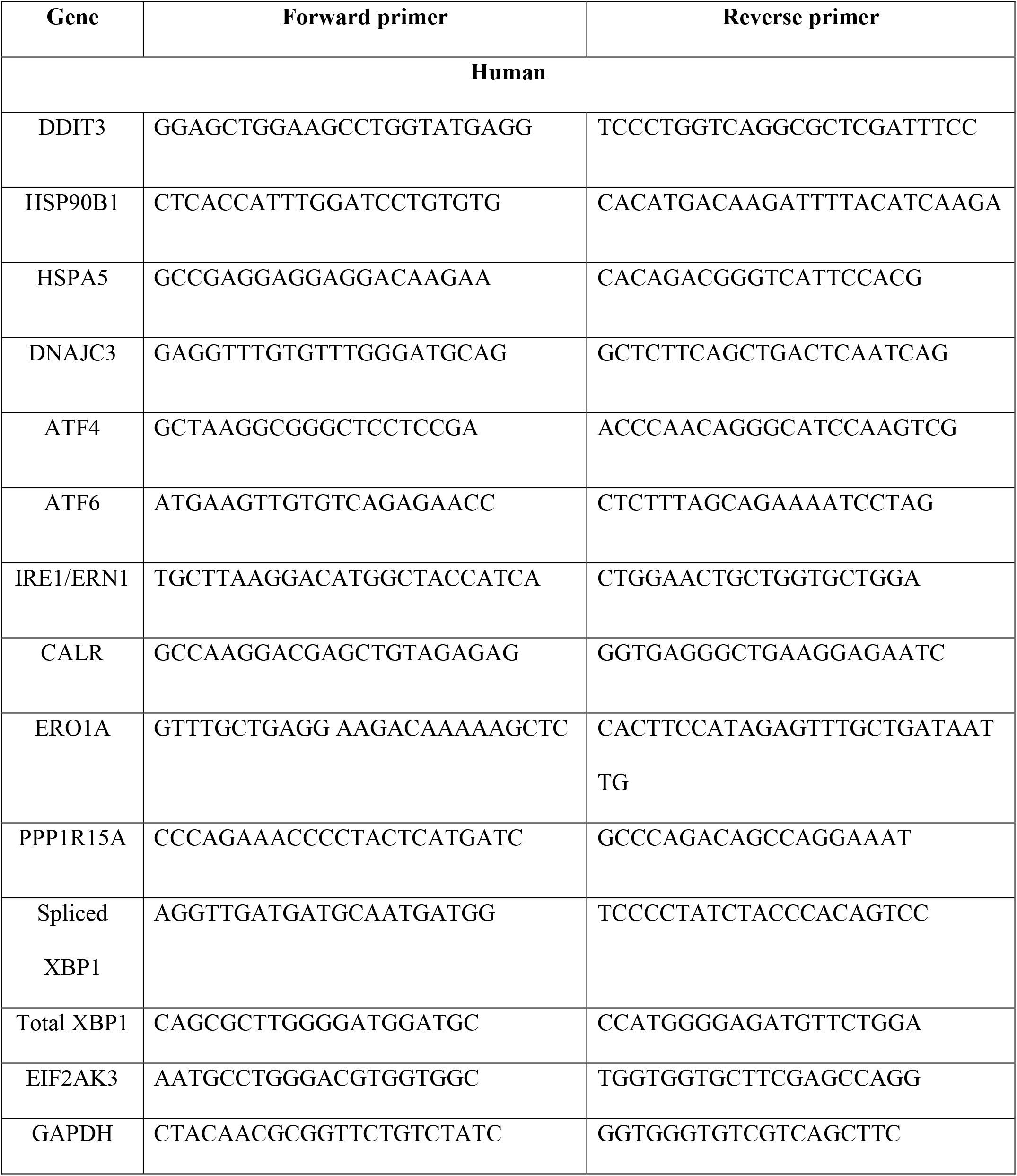
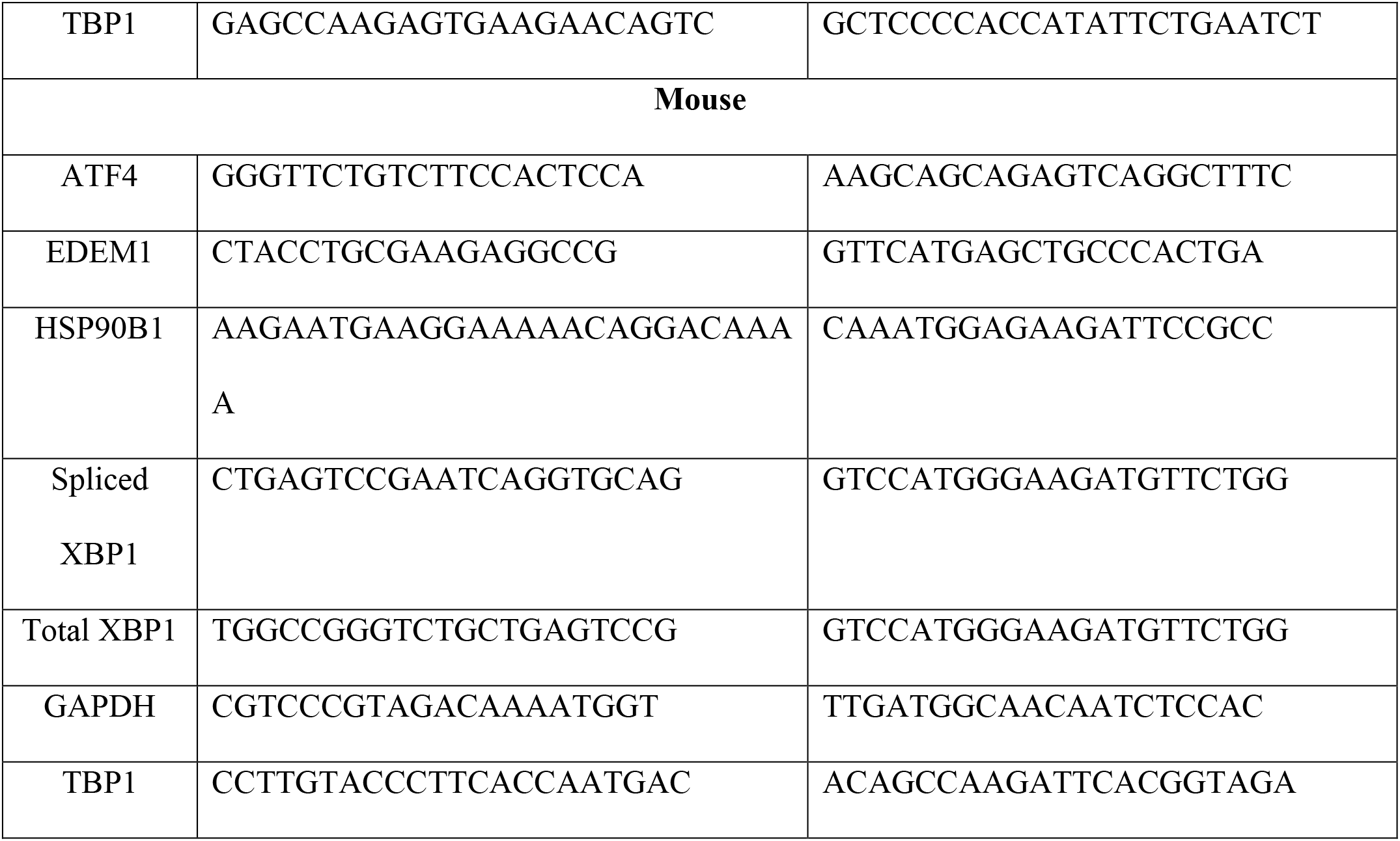
Primer combinations used for quantitative RT-PCR.

